# Resting-State fMRI Dynamics and Null Models: Perspectives, Sampling Variability, and Simulations

**DOI:** 10.1101/153411

**Authors:** Robyn Miller, Anees Abrol, Tulay Adali, Yuri Levin-Schwarz, Vince Calhoun

## Abstract

Studies of resting state functional MRI (rs-fRMI) are increasingly focused on “dynamics”, or on those properties of brain activation that manifest and vary on timescales shorter than the scan’s full duration. This shift in focus has led to a flurry of interest in developing hypothesis testing frameworks and null models applicable to the dynamical setting. Thus far however, these efforts have been weakened by a number of crucial shortcomings that are outlined and discussed in this short paper. We focus here on aspects of recently proposed null models that, we argue, are poorly formulated relative to the hypotheses they are designed to test, i.e. their potential role in separating functionally relevant BOLD signal dynamics from noise or intermittent background and maintenance type processes is limited by factors that are fundamental rather than merely quantitative or parametric. In this short position paper, we emphasize that (1) serious care must be exercised in building null models for rs-fMRI dynamics from distributionally stationary univariate or multivariate timeseries, i.e. timeseries whose values are each independently drawn from one pre-specified probability distribution; and (2) measures such as kurtosis that quantify over-concentration of observed values in the far tails of some reference distribution may not be particularly suitable for capturing signal features most plausibly contributing to functionally relevant brain dynamics. Other metrics targeted, for example, at capturing the epochal temporal variation that contributes heavily to dynamic functional connectivity estimates and is and often taken as a signature of brain responsiveness to stimuli or experimental tasks, could play a more scientifically clarifying role. As we learn more about the phenomenon of functionally relevant brain dynamics and its imaging correlates, scientifically meaningful null hypotheses and well-tuned null models will naturally emerge. We also revisit the important concept of distributional stationarity, discuss how it manifests within realizations versus across multiple realizations, and provide guidance on the benefits and limitations of employing this type of stationarity in modeling the absence or functionally relevant temporal dynamics in resting state fMRI. We hope that the discussions herein are useful, and promote thoughtful consideration of these important issues.

## 1. Introduction

Studies of blood oxygenation-level dependent (BOLD) resting state functional magnetic resonance imaging (rs-fMRI) have been increasingly focused on properties of functional activation that manifest and vary on timescales shorter than the full duration of the scan 1. Such an approach is a more natural way to analyze such data as we know the brain is a highly dynamic organ. In resting-state studies, we do not have the benefit of external indicators of behavior other than the narrow case of subject wakefulness, which can be studied by using simultaneous electroencephalogram (EEG) recordings^2–4^ In the case of resting data, we are seeking evidence in the scan itself of shifts in a subject’s cognitive focus, emotional state, attention or consciousness level: e.g., we are attempting to locate the temporal and correlative signatures of complex internal (“ecologically authentic”) sequences of mental tasks. The growing focus on shorter timescale analysis and examining these functionally relevant brain dynamics (FRBD) in rs-fMRI has spawned a number of efforts^1,5–7^ to identify variations (univariate, multivariate and relational/correlative) in network behavior that arise from reconfigurations of the subject’s *cognitive, attentional, sensory, emotional* (CASE) states. These shifts can be conscious or non-conscious, and targets of investigation can be expanded to include temporal fluctuations that shape and maintain crucial neural circuitry that indirectly supports effective brain function.

Testing the statistical significance of the dynamic (i.e. time-varying) measures capturing these CASE shifts in such shorter timescale analyses assumes great importance because of the noisy (inherent physiological and artefactual confounds) nature of BOLD rs-fMRI data. While it would be highly useful to replicate the behavior of “noiseless” BOLD data to construct “appropriate” null simulations, the absence of a baseline, i.e. ground truth for resting state, makes this step extremely challenging. A null distribution of the test-statistic capturing the phenomenon of interest must therefore be approximated using multiple, independent “surrogate” realizations of the empirical data. The surrogate data realizations ideally retain all statistical properties of the empirical data other than the phenomenon of interest, and hence give a meaningful null to validate statistical significance of the metric capturing the phenomenon of interest. Notably, non-parametric null models based on phase randomization (PR) and/or vector auto-regression (VAR) approaches have been widely used to seek evidence for presence of non-stationarities (discussed in section 2.1). These models allow us to comment on the Gaussian, stationary, and linear properties of the studied data. The work on null models and hypothesis testing (e.g.,^8–14^) frameworks for shorter timescale analysis of network behavior and dynamics has produced some preliminary (although possibly contrasting) insights but limitations remain. One major limitation of using these (phase randomized or vector autoregressive) non-parametric null models as also pointed in a few of these studies^13–14^ is that they can also be rejected due to presence of non-linearities. Because of this, these approaches in their current form, do not allow us to conclude the presence/absence of non-stationarities in the case of rejection of these nulls. More importantly, the fact that values within a signal are not inconsistent with values drawn from some fixed pre-specified distribution, i.e. that the signal is not provably distributionally nonstationary does not rule out the presence of FRBD^14^ Given the dynamic nature of the human brain, a more interesting hypotheses about fMRI dynamics would not focus on *whether* they exist, but rather on how they might manifest differentially over different temporal, spatial, and functional scales.

The best we can hope for right now is for researchers to focus on what signal properties they are seeking to quantify, and why they believe that a strong presence of these properties should be taken as evidence of FRBD (or of nuisance factors that are not easily separable from FRBD at current levels of measurement resolution). It can be argued that analyzing univariate or multivariate timeseries variations that are most plausibly connected with actual shifts in mental functioning (e.g. temporal epochs exhibiting profound changes in magnitudes and spectrum) will ultimately yield more scientifically clarifying information about observed resting state brain dynamics than approaches based on quantifying the improbability of observed values with respect to some fixed reference distribution. As we will be discussing in this paper, the presence of spectrally distinguishable temporal epochs at both univariate and multivariate scales (where they play a larger role in connectivity) is one potential form of evidence for rs-FRBD that might warrant more attention. There are many features and timescales over which functionally relevant temporal variations might manifest, and most prospective metrics will present some combination of over-sensitivity to irrelevant features and blindness to important features and/or timescales. Development of valid metrics for functionally relevant brain dynamics presents thus presents a major challenge in the field (as also discussed in^10^). Such a metric would essentially, with some degree of specificity, rise in the presence of those univariate or multivariate timeseries variations that are most plausibly connected with actual shifts in mental functioning. The factors that obstruct development of powerful, valid metrics also present serious challenges to the development of valid, scientifically useful null models. In studies where the natural null hypothesis is effectively that “measured dynamic signal properties do not reflect functionally relevant dynamic neural processes,” an appropriate null space of network timecourses would have to lack variations consistent with actual shifts in mental functioning, for example, (1) there must be no task, experimental or ecological condition whose signature presents as a type of epochal variation^1^ generically observable in this null space, and (2) the timeseries features that occur most rarely in this space must be exactly those that are most strongly consistent with a brain undergoing shifts in CASE state. Another, more general, hidden risk when using multi-parameter simulation-based null models is ensuring that the resulting distributions of key test statistics are not influenced by auxiliary model parameters unrelated to those explicitly connected with the null hypothesis.

**Figure 1.**
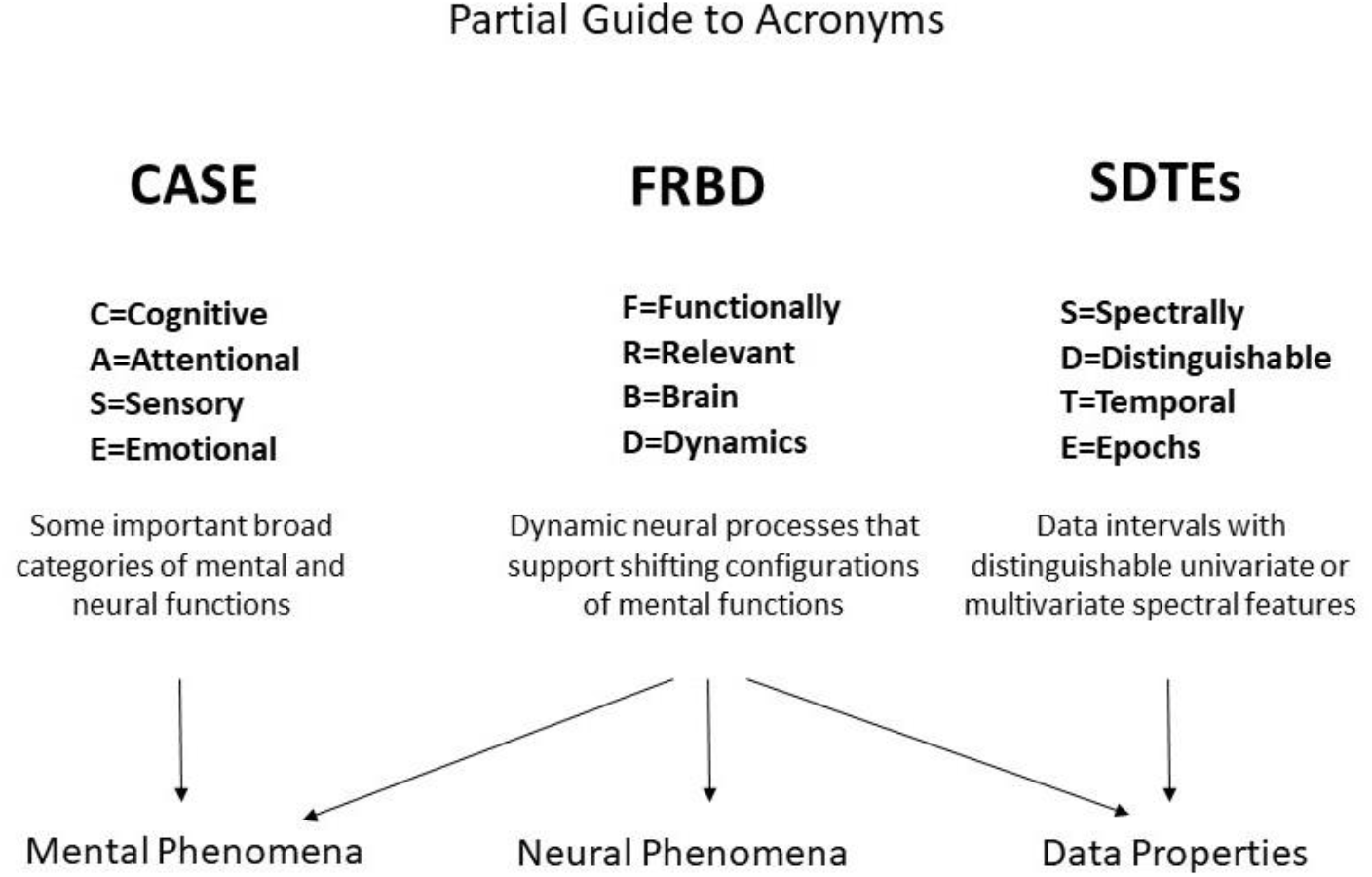
Diagrammatic summary of several key acronyms in use throughout this paper.

In this short position paper, we further detail these points and would like to address them in particular as they relate to the null model and test metric used in a recent paper^9^ Our concerns are applicable well beyond the scope of^9^ but this recent paper motivates us to discuss some of the inherent challenges of dynamic fMRI analyses, and helps illustrate the key pitfalls of employing naive null models in efforts to detect the presence of ubiquitous, complex and poorly understood phenomena such as functionally-relevant brain dynamics.

As this position paper was written in partial dialogue with a recent published paper^9^addressing the validity of dynamic functional network connectivity, we provide a very brief summary of some salient details from that paper to facilitate the reader’s engagement with the discussion that follows. Using a null space of windowed connectivity matrices computed from spectrally and covariance constrained Gaussians (SCC Gaussians), the authors employ multivariate kurtosis as a test statistic to test the null hypothesis of “no dynamics”. Among other findings, they report (1) a strong relationship between motion (mean frame displacement) and kurtosis; and (2) that the clusters formed by time-varying connectivity matrices from the null space are nearly indistinguishable from those computed from empirical data. Also, while the focus of^9^ was predominantly on connectivity dynamics, connectivity-based findings do carry much broader implications about the validity of shorter timescale dynamics of fMRI voxel and network timecourses as well (if connectivity is not dynamically changing on some timescale, then the underlying univariate signals are not dynamically changing in any ways that validly affect their measured correlative relationships etc.).

We hope our discussion clarifies the implications of^9^, while also conveying the challenges of the project its authors undertook. It must also be noted that for remainder of the manuscript we shall be focusing on discussing appropriateness of null models in the context of epochal temporal variations (key contributors to estimated FC measures and potential signatures of FRBD) as the specific phenomenon of interest, and not in context of testing for scan-length statistical stationarity (unless otherwise specified).

## 2. Modeling Functionally Relevant Brain Dynamics vs. Modeling Their Absence (Data Models vs. Null Models)

The relationship between well-constructed models of observed data and null models for a given phenomenon depends largely on the relationship between observed/observable data and the phenomenon being investigated. In situations such as those presented by the study of resting-state functionally relevant brain dynamics (rs-FRBD) using human subject fMRI data, where the null hypothesis is that a certain phenomenon is not present but the empirical data being modeled (in this case real fs-fMRI data) happens to be data in which this phenomenon is continually present, then the goals of building a null model for the phenomenon and accurately modeling the data can diverge significantly. Though there are many cases where highly accurate models of the data are also appropriate models of a setting in which the null hypothesis applies, e.g., they are also good null models. In the fMRI setting, since we tend to correct for motion artifacts, then a good model of this data that might, for example, be useful for identifying new scans that should be examined for motion contamination.

As discussed earlier, one of the challenges in the field of brain dynamics is that it is difficult to generate a true/valid null model as the phenomena of interest are rather poorly defined. Thus, modeling this data with high fidelity is not going to be the same as producing a null model for brain dynamics. In fact, a valid null simulation model of multivariate signals lacking features corresponding to rs-FRBD would by necessity diverge from actual scans observed in living people. Depending on what turn out to be the most reliable timeseries indicators of CASE-driven brain dynamics, it is possible that there could be a valid null model that exhibits some similarities with the observed data (or output from good models of that data). However, a model built on the null hypothesis of no brain dynamics would by necessity only rarely create realizations that look similar to real rs-fMRI multivariate timeseries.

### 2.1. Statistical Stationarity, Gaussianity and rs-Brain Dynamics

Statistical or distributional stationarity is defined through the invariance of its joint probabilistic distribution across any number of samples, to any time shift, and as such, with real data, it can be only inferred using multiple realizations of a given stochastic process. A process is stationary if the statistical characterization computed over *N* realizations of any *k*-tuple of timepoints of length *j*{*t*_*j*_1__,…,*t_j_k__*} and their *τ*-translates {*t*_*j*_1__ + *τ*, …,*t_j_k__* + *τ*) converge to identical values as *N* → ∞. A practical way to infer stationarity is by estimating a finite set of moments as they are easier to compute than a full probability distribution. Of course, it is important to note that any definition of stationarity depends on the interval over which it is evaluated. For a given interval over which stationarity holds it is quite possible for strong nonstationarities to manifest over shorter sub-’intervals as we demonstrate next. Matching white Gaussian noise to a template spectrum actually produces every possible signal with the given time-averaged spectral content given by the target. Statistical stationarity without other explicit constraints on the process does not imply that individual realizations of the univariate or multivariate timeseries (e.g., for fMRI these are individual subjects) are not featuring pronounced temporal epochs (see **Figure 2**). Even white Gaussian noise, for example, matched to some empirically-valid band-limited spectrum – a common statistical tactic that was also a step in the null model proposed in^9^ – can be markedly epochal within typical individual realizations. This general phenomenon is mostly clearly visually evident in cases like the one presented in **Figure 3** where Gaussian white noise is spectrally matched to a narrow-band spectral template.

**Figure 2.**
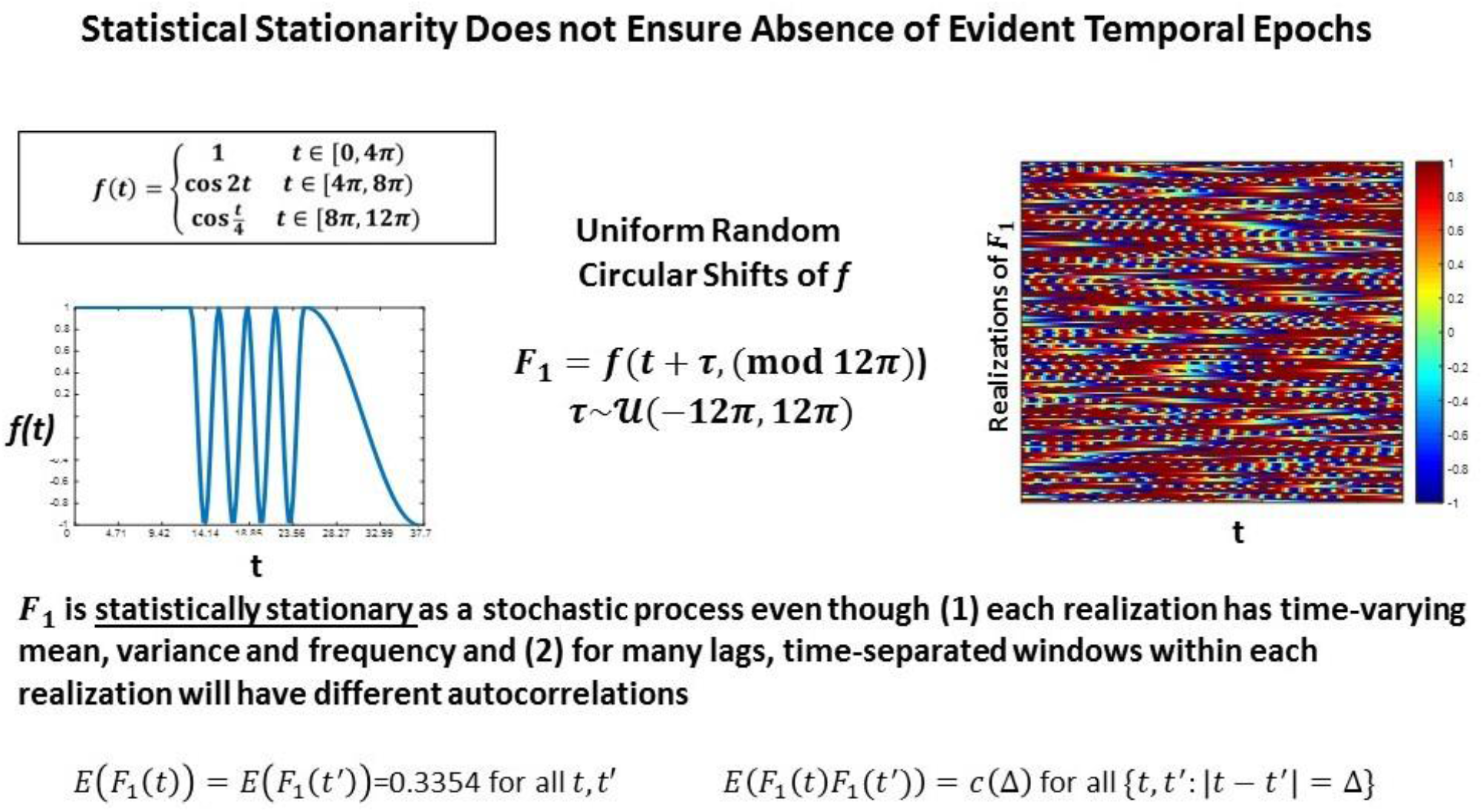
The function f is a highly stylized example of a signal with distinguishable temporal epochs. The first, second, and final third have different means, variances and characteristic frequencies. The stochastic process F_1_, however, whose realizations are obtained through uniform random circular shifts of f, is statistically stationary in that that the statistical summaries assessed at distinct timepoints over large numbers of realizations are the same. All realizations have spectral and epochally clear variations, which would be reflective of FRBD, but as a stochastic process the collection of phase-shifted versions of f are statistically stationary.

**Figure 3.**
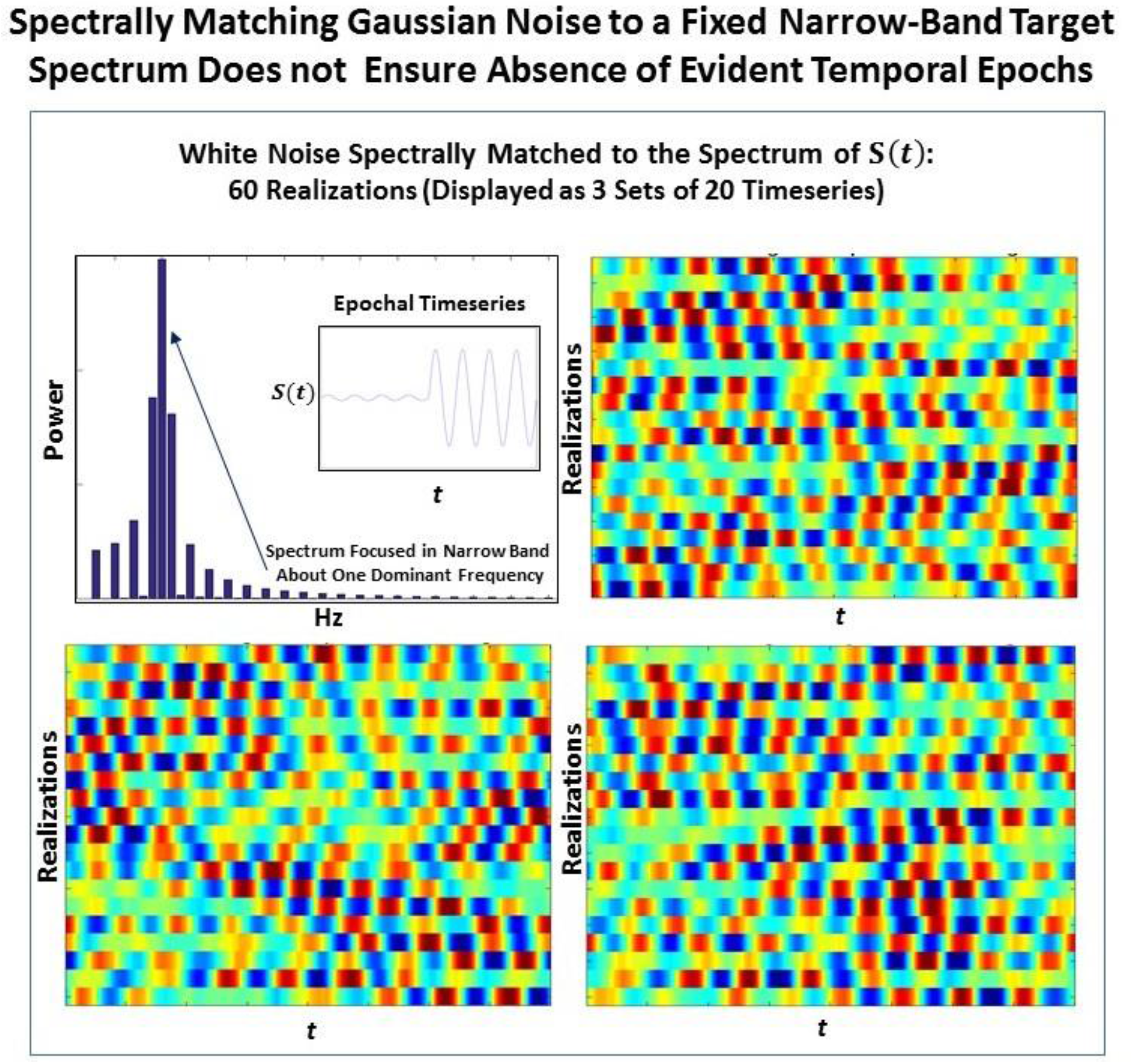
The signal S(t) consisting of a low-amplitude 0.08Hz segment followed by a high-amplitude 0.08 Hz segment is one manifestation of a signal with a narrow-band spectrum (shown top left) focused at 0.08 Hz. Matching 60 timeseries of Gaussian white noise to this spectrum yields (in sets of 20) the timeseries shown in the other three panels of this figure. It is evident that spectral-matching of Gaussian noise to a generic template spectrum can contain discernible temporal epochs with any given realization. Thus, such a model cannot be a good null model for dynamic connectivity as it will contain the very dynamics we are interested in studying.

The simulated null model of^9^ is built from spectrally and covariance constrained multivariate Gaussian processes (SCC Gaussians). The use of an SCC Gaussian timeseries as a null model for functionally-relevant brain dynamics (FRBD) rests on an implicit assumption that (in the case of a 95% confidence level) fewer than 5% of the signals in this space exhibit characteristics consistent with FRBD. The use of fMRI as an imaging modality also rests on an implicit assumption that some significant proportion of the data being recorded reflects functionally relevant brain dynamics. It would be hard, for example, to justify studying even scan-length (static) spectral and relational/connectivity characteristics of timeseries whose fluctuations are believed to be largely artifactual. The functioning human brain during any state of wakefulness is (hopefully inarguably) continually engaged in myriad temporally-varying combinations of cognitive, sensorimotor, attentional, emotional, planning, imagination and memory-related tasks. Many of these functions are in use even during sleep. Thus, any null model of multivariate timeseries whose characteristics are highly consistent with empirically observed fMRI-based brain measurements has little utility (notably in the context of functionally relevant brain dynamics), since the phenomenon that it is testing for is ubiquitous rather than rare. The space of SCC multivariate Gaussians replicates real fMRI network timeseries with sufficient fidelity to induce broad consistency in measurable characteristics between the simulated data and the empirical data it was modeled upon^14^ Moreover, there is no *a priori* reason to believe that aberrant or “tail” phenomena in this space should be more strongly associated with functionally-relevant brain dynamics than with measurement noise, motion or other artifacts, e.g., the sort of features that might warrant examining a scan for possible removal rather than positioning it as an exemplar of functionally-relevant resting state brain dynamics.

### 2.2. Multivariate Signals Lacking Plausible Markers of rs-Brain Dynamics (Valid Null Models for Brain Dynamics)

The development of valid null models for rs-FRBD is substantially hindered by a dearth of fMRI recordings from living subjects under conditions that all but preclude the ongoing CASE shifts unavoidably present even in sleeping or mentally impaired subjects. Resting state fMRI data is generally recorded under conditions in which functionally relevant brain dynamics ought to be continually present. Thus, the measurable features of empirically observed multivariate fMRI network timeseries are intractably “contaminated” from the standpoint of parameterizing a null space in which signal properties reflecting rs-FRBD are ensured to be rare.

Another challenge for hypothesis testing of rs-FRBD resides in identifying quantifiable signal features for which every upward increment of the associated measure unambiguously yields stronger evidence for the presence of rs-FRBD. Without this property, observations from the distributional tails of the measure are simply improbable, but not necessarily in ways that relate to FRBD. Thus, the first-layer of challenges is posed by our limited understanding of the signal properties whose variation through time reflect some shift in one of the brain’s myriad comingling functions. These are often amplified by non-monotonic relationships between those properties and the neural functions they putatively reflect. One example of such a property is kurtosis (see Technical Supplement), a higher-order statistical moment that has been employed^9^ to gauge the (presumably function-relevant) “dynamic-ness” of simulated and empirical fMRI signals. Univariate kurtosis captures the “peakedness” of a unimodal distribution; itrises with the number and magnitude of observations in a sample that would be outliers if the underlying process was stationary Gaussian, i.e., a Gaussian with constant mean and variance. Modestly elevated kurtosis might well reflect some unusually strong or active brain dynamics – this would have to be demonstrated, but is not implausible. However, extremely large kurtosis values occur when a sample contains numerous wildly large (in magnitude) observations. A brain recording with these characteristics is more likely to suggest noise, motion or other nuisance factors than anything connected with actual rs-FRBD occurring during the scan (see **Figure 5**). More generally though, the effort to statistically validate the presence of a phenomenon that is almost axiomatically continual in any valid recording seems misguided. The more interesting hypotheses about fMRI dynamics would not focus on *whether* they exist, but rather on how they might manifest differentially over different timescales, spatial scales and functional scales.

## 3. Measuring Functionally Relevant Brain Dynamics vs. Identifying Outlying Observations

A valid metric of brain dynamics should rise monotonically with the strength of the signal features that are, at our current level of scientific knowledge, widely believed to have associations with CASE or task-driven shifts in brain function. The metric should also be as blind as possible to signal features believed to represent nuisance factors. The problem of quantifying signal features that have a high likelihood of representing evolving CASE states is admittedly very difficult. Every procedure will be biased by assumptions whose validity cannot be ensured based the current state of knowledge. The best we can hope for right now is for researchers to be clear about what signal properties they are seeking to quantify, and why they believe that a strong presence of these properties should be taken as evidence of FRBD (or of nuisance factors that are not easily separable from FRBD at current levels of measurement resolution).

We show in this section that a measure based on kurtosis, while sensitive to outliers, is not an ideal metric to capture brain dynamics and it is quite easy to show that kurtosis can be more sensitive to very rare outliers than it will be to more prevalent FRBD. We show in both stylized examples but also in real data that kurtosis preferentially captures signal features likely to arise from measurement disruptions (e.g., motion), while suppressing evidence of more extended spectral epochs within network timeseries. We also propose a new metric, **Φ**, which we believe shows some of the desired properties.

### 3.1. Univariate and Multivariate Kurtosis Under Stationary Gaussianity

Univariate excess or normalized kurtosis, the fourth statistical moment rescaled by squared variance and centered by subtraction of 3 has a well-understood distribution under the Gaussianity assumption that is applicable to^9^ and here. Under this assumption, there is a closed-form transformation (dependent on the sample size, *n*) that converts observed excess kurtosis 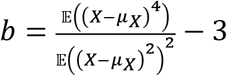 into an unbiased estimator of true kurtosis, 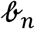 (see Technical Supplement) that is distributed as a standard normal 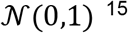 and hence now has altered limits. The use of 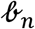 enables statistical evaluation of departures from stationary Gaussianity. When we refer to values of univariate kurtosis, these are values of the unbiased estimator 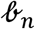. Samples that yield elevated values of 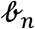 (say, 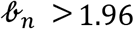, so that *p* < 0.05) contain high-magnitude observations that are too numerous and/or too extreme for the sample to have even a 5% chance of having been generated by a stationary Gaussian process.

There is a similar transform for Mardia’s multivariate kurtosis (m.v. kurtosis), with a similar interpretation. The unbiased estimator 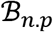 (dependent on sample size, *n*, and vector length, *p*, see Technical Supplement) for Mardia’s multivariate kurtosis^16^ is:

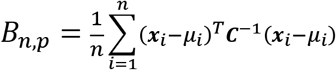

where ***C***^−1^ is the *n* × *n* inverse covariance matrix of the time-indexed *p*-vectors {***x***_1_, ***x***_2_,…,***x***_*n*_}.

### 3.2. Wavelet-Based Metric of Spectrally-Distinguishable Temporal Epochs

We briefly introduce a novel metric 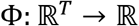 that explicitly captures the presence of spectrally distinguishable temporal epochs in a timeseries (see **Figure 4** and the Technical Supplement). The metric has a natural multivariate extension 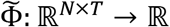 that evaluates spectrally distinguishable temporal epochs in multivariate timeseries. Φ is not a primary focus of this short paper, but it plays a role in the discussion that follows because it provides a more targeted measurement than, for example, kurtosis, of timeseries characteristics that could form FRBD.

**Figure 4.**
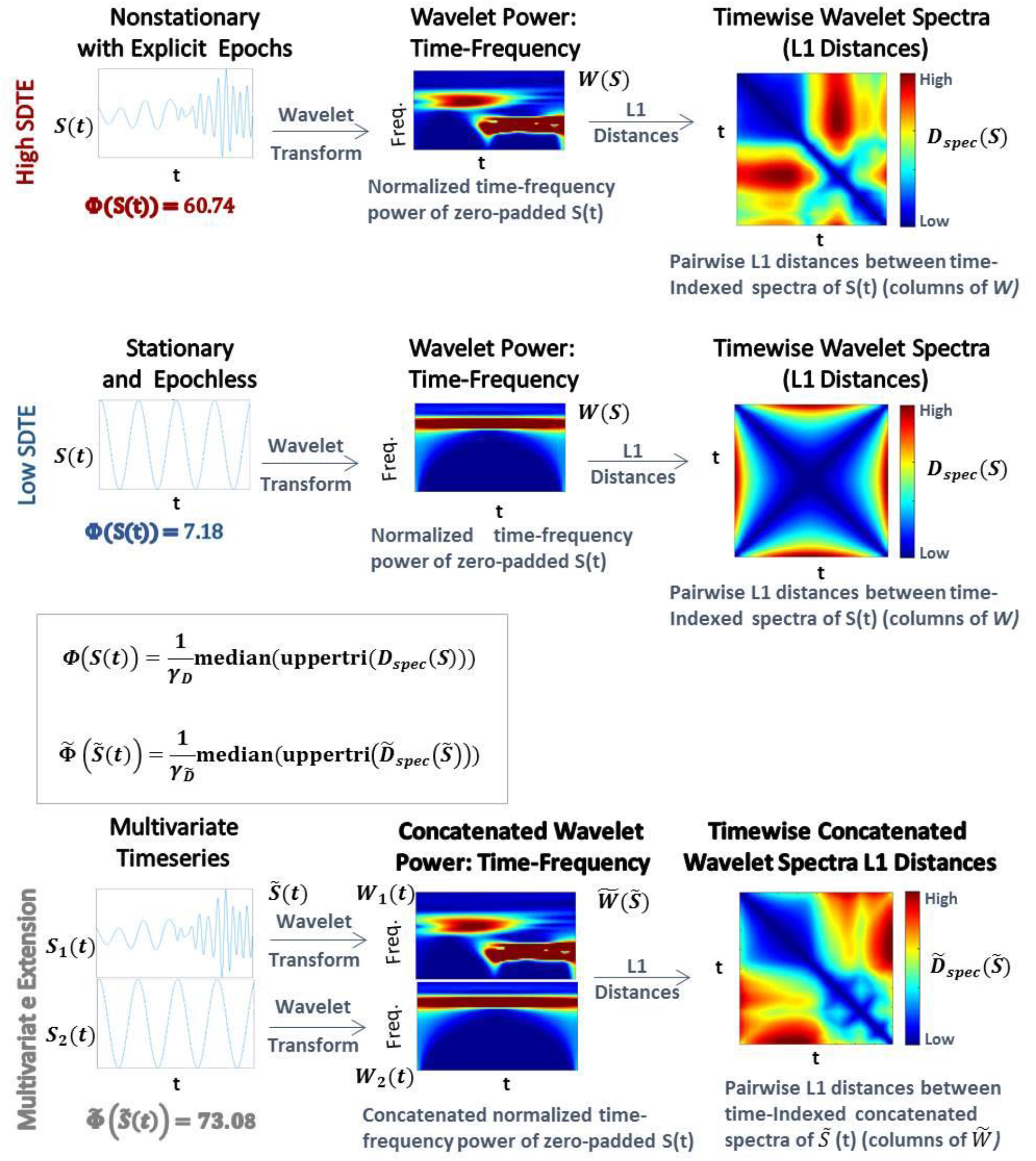
The metric Φ is intended to capture spectral nonstationarities, or spectrally distinguishable temporal epochs (SDTEs), in univariate (top) and multivariate (bottom) timeseries. The univariate version uses wavelets to capture temporally-localized spectral information, yielding a set of time-indexed spectra (top middle), which we normalize to have total power equal to the product of the number of frequencies and the number of timepoints. The spectrum at each timepoint carries modest edge effects, which are more pronounced near the signal boundaries and for slower frequencies captured by wavelets that do not taper as much near the boundaries. We then compute pairwise L1 distances between the time-indexed spectra (top right) and compute the median off-diagonal value of the resulting T × T matrix (scaled by a scaling factor γ; see Technical Supplement for details). The multivariate extension 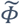 of Φ concatenates the time-frequency spectra of all univariate constituents along the frequency dimension (bottom middle), leaving the time dimension unaltered. In this case, we compute the median off-diagonal elements of the T ×T matrix of pairwise differences between time-indexed concatenated spectra (bottom right). The multivariate metric is higher when constituent univariate time-frequency spectra exhibit their largest within-signal spectral differences in mutually distinct temporal intervals. The case shown here does not illustrate the role of differential intervals of SDTEs among constituent timeseries. The multivariate extension of this metric was successful in capturing slight changes in frequencies in several consecutive short term epochs too (Supplementary Figure 1). Finally, in all cases, the results were highly similar for both L1 (Manhattan) and L2 (Eucledian) distances (multivariate ϕ outperformed multivariate kurtosis for both distance measures).

### 3.3. Epochal Stationarity and Kurtosis

The presence of spectrally distinguishable temporal epochs across realizations, i.e., multiple subjects’ connectivity characteristics in individual or multivariate network TCs is one reasonable potential form of evidence for rs-FRBD. Although it is also possible that this type of phenomenon could arise from nuisance factors, epochal behavior has structure that makes it less likely to be sourced dominantly in nuisance factors such as motion, measurement noise or physiological rhythms. Kurtosis, which can help identify the presence of outliers in Gaussian data, has been proposed as a metric to detect FRBD, however kurtosis is highly susceptible to unstructured amplitude variations. Moreover, as an outlier metric, kurtosis has greater sensitivity to sharp, transient, high-amplitude anomalous intervals than to signals with amplitude and frequency variation on more functionally relevant timescales (see **Figure 5**). In fact, the properties leading to elevated kurtosis are sometimes more present in an epochally stationary signal than in an epochally nonstationary signal (see **Figure 6**), i.e., one that is stationary except within a given duration. This is not to say that measuring epochal nonstationarity is straightforward. There are many features and timescales on which the nonstationarity might be exhibited, and most prospective metrics will present some combination of oversensitivity to irrelevant features and blindness to important features and/or timescales. We are currently working on a flexible, tunable approach to capturing the kind of epochally structured frequency domain variation that promises to provide valid evidence for brain dynamics after careful evaluation of sensitivity to nuisance factors.

**Figure 5.**
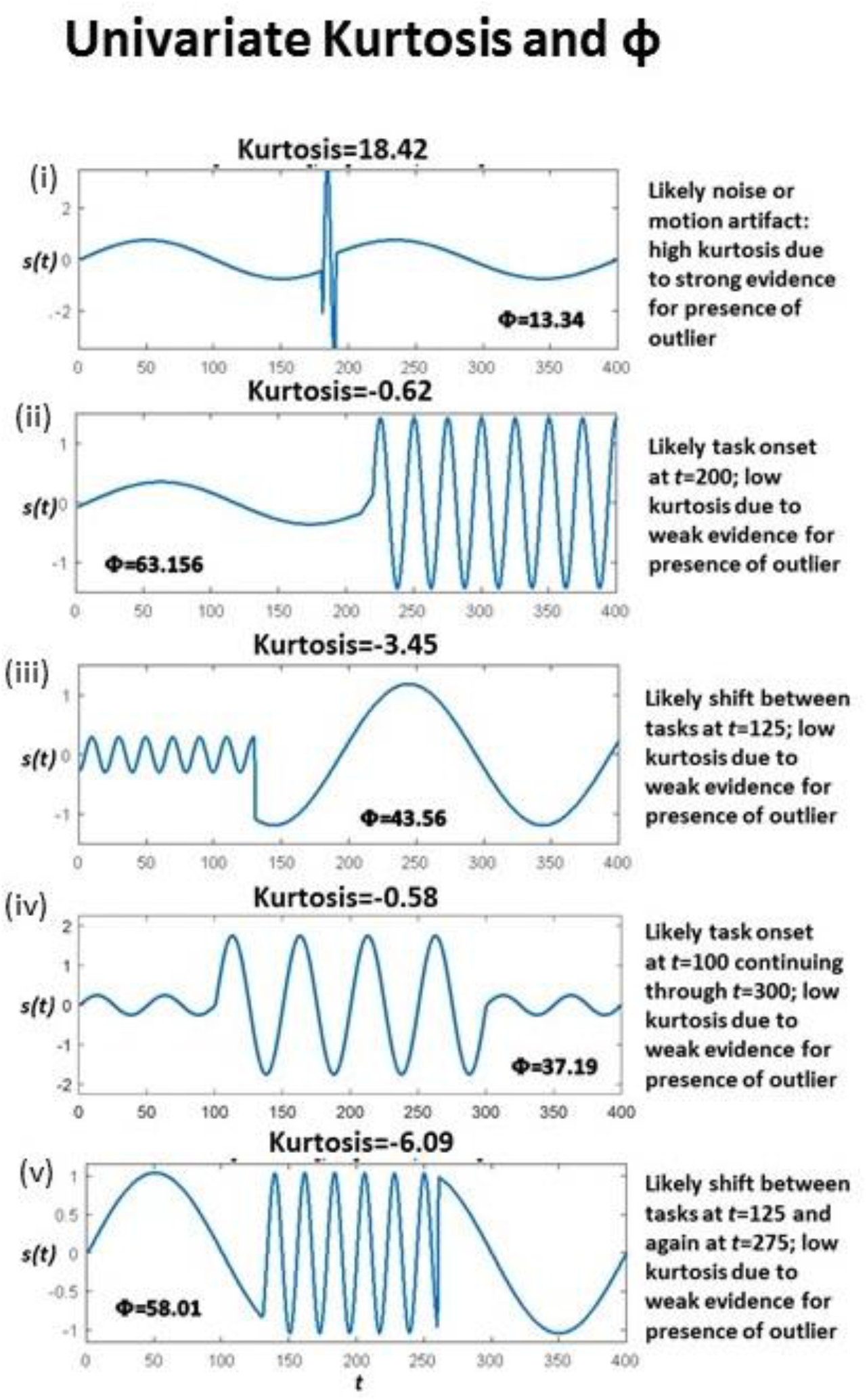
(i) Highest kurtosis applies in a signal with a transient high-amplitude high-frequency interval, more consistent with a motion or noise than with shifting CASE states; **Φ** is extremely low for this seeming artifact; (ii-iv) Much lower kurtosis in stylized signals with highly distinguishable temporal epochs are more consistent with FRBD; much higher **Φ** in these strong FRBD candidates; (v) Lowest kurtosis in stylized signal with very strong and distinguishable temporal epochs; much higher **Φ** for this strong FRDB candidate. Positive univariate excess kurtosis indicates super-Gaussianity (and is not particularly useful for indicating functionally relevant dynamics). As such, kurtosis is high when there are a larger number of high-amplitude observations than should arise under the assumption of Gaussianity. Univariate kurtosis (transformed via equation (1) in the Technical Supplement) to distribute, assuming Gaussianity, as a standard normal random variable) is negative on each of the stylized examples (ii)-(v) that exhibit distinguishable temporal epochs consistent with functionally relevant brain dynamics. It is very large and positive only in the example containing a single high-amplitude, high-frequency “spike” (i). The behavior that appears in the upper tail of the kurtosis distribution is more consistent with motion artifacts or measurement error than anything previous imaging or EEG studies have found to be associated with experimental tasks. The metric, Φ, introduced in this work, is at least three times larger for examples (ii)-(v) that exhibit distinguishable temporal epochs consistent with functionally relevant brain dynamics than for the case (i) that features a single high-frequency high-amplitude “spike”embedded in an otherwise spectrally epochless signal. As such Φ exhibits the behavior we would expect, whereas kurtosis is not particularly useful for detecting behavior consistent with relevant brain dynamics for the examples shown above. Additionally, Φ successfully captured small changes in frequencies in several consecutive short term epochs as well (Supplementary Figure 2).

**Figure 6.**
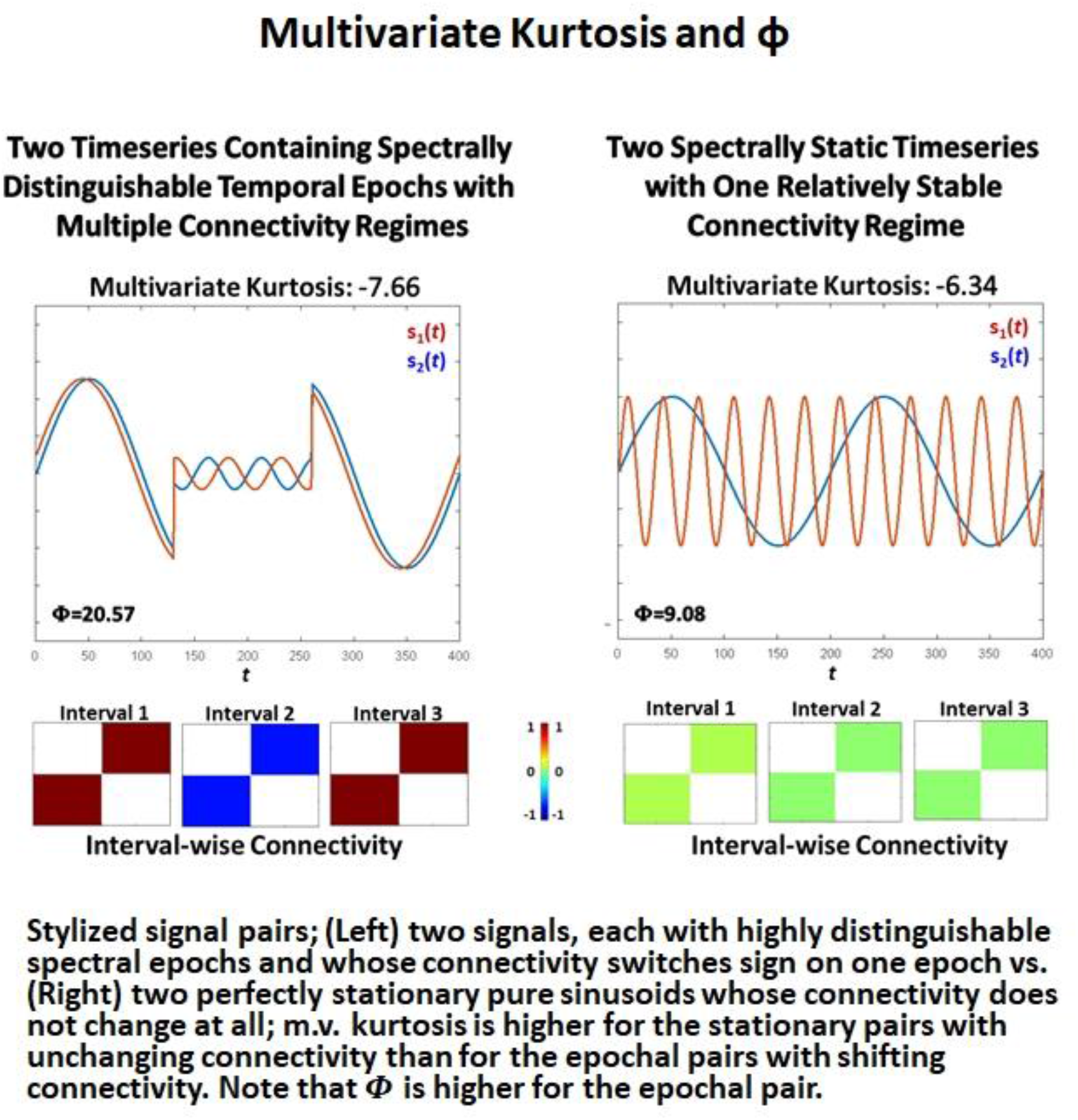
Stylized signal pairs; (Left) two signals, each with highly distinguishable spectral epochs and whose connectivity switches sign on one epoch vs. (Right) two perfectly stationary pure sinusoids whose connectivity does not change at all; m.v. kurtosis is higher for the stationary pairs with unchanging connectivity than for the pairs with shifting connectivity. **Note that Φ is higher for the pair the includes** epochs. Excess multivariate kurtosis indicates multivariate super-Gaussianity (and is not particularly useful Stylized signal pairs; (Left) two signals, each with highly distinguishable spectral epochs and whose connectivity switches sign on one epoch vs. (Right) two perfectly stationary pure sinusoids whose connectivity does not change at all; m.v. kurtosis is higher for the stationary pairs with unchanging connectivity than for the epochal pairs with shifting connectivity. Note that Φ is higher for the epochal pair. for indicating functionally relevant dynamics). Multivariate kurtosis (transformed via equation (4)) from the Technical Supplement) is assumed to be distributed, assuming multivariate Gaussianity on the part of the random vector, as a multivariate standard normal random variable) on both stylized multivariate examples. The first (left) features two signals that each exhibit highly distinguishable spectral epochs and whose correlative behavior is also dynamic: they are perfectly correlated, then perfectly anti-correlated, then again perfectly correlated. This is a very dynamic context but not only presents negative multivariate kurtosis, its kurtosis value is even more negative than the second example (right) that features two spectrally unchanging signals whose mutual correlations are consistently zero. The multivariate measure Φ is twice as large in the dynamic example (left) compared to the static example (right).

### 3.4. Empirical Data and Simulation Regimes

A set of network timecourses from a clinical rs-fMRI study on which dynamic functional network connectivity (FNC) results have already been published^17^, and five simulation regimes modeled on that data are employed to explore and illustrate the role of Gaussianity and statistical stationarity as well as, spectral and covariance stationarity in modeling rs-FRBD (and/or its absence). It is important to note that we refer to stationarity in the true sense as statistical (non)stationarity in what follows. We use the terms “spectral (non)stationarity” and “covariance (non)stationarity” to refer to other definitions used including^9^ that invoke the concept of (non)stationarity through the analysis of a single realization either in the spectral domain (for the former definition) or using covariance function (for the latter definition). The approaches are lightly outlined here, with more details provided the Technical Supplement.

#### Real Data

We used previously published^17^ network timecourse data from a large multisite clinical resting-state fMRI study. These timecourses (314 subjects, 47 networks, 158 timepoints), subsequently filtered for frequencies at most 0.08 Hz and z-scored, are referred to below as “Real Data”^17^ (see **Figure 7**; top left). The average power in each frequency bin in [0.003,0.08] Hz for all network TCs for all subjects is denoted 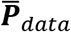. The average cross-network covariance matrix for all subjects is denoted 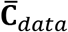. Every simulation regime described below consists of 1000 simulated subjects, each characterized by a set of 47, length-158 timeseries.

**Figure 7.**
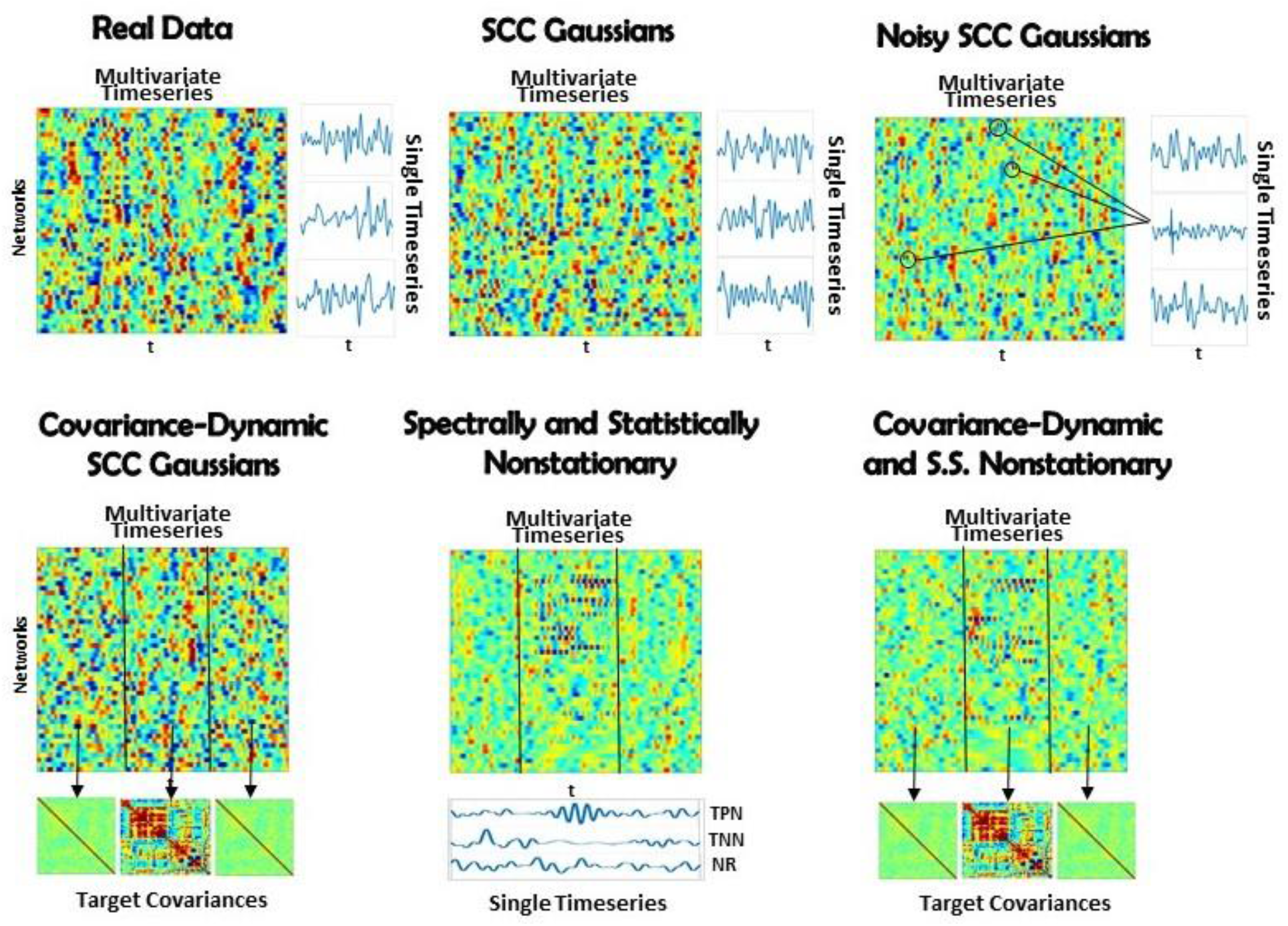
(Top Left) Real Data: Multivariate timeseries consisting of network timecourses from an actual rs-fMRI subject^17^ (3 single network examples displayed immediately to the right); (Top Middle) SCC Gaussians: Multivariate timeseries of SCC Gaussians modeled on mean spectral content and mean mutual covariance of empirical network timecourses from the study shown at the top left.; (Top Right) Noisy SCC Gaussians: The same as SCC Gaussians but with between 3 and 15 high-amplitude, high-frequency “spikes” interspersed, examples circled. No more than one spike is inserted in any individual timecourse and at most 15 of the 47 timecourses contain spikes. (Bottom Left) Covariance-Dynamic SCC Gaussians: This regime starts like the SCC Gaussians, in that the timeseries are spectrally matched to the mean spectral content of the target empirical dataset. However, the next stage involves covariance matching the middle interval to the mean mutual covariance of the empirical networks, while the first and final intervals are matched to a very weakly connected covariance structure. (Bottom Middle) Spectrally and Statistically Nonstationary: This regime also starts like the SCC Gaussians, in that the timeseries are spectrally and covariance matched to the mean spectral content and covariance structure of the target empirical dataset. Here though there is a middle window in which a subset of TPNs exhibits high-amplitude, high frequency behavior, a subset of TNNs exhibits low-amplitude, low-frequency behavior and most networks are NR to the stimulus. (Bottom Right) Covariance Dynamic and SS Nonstationary: This regime starts like the Covariance Dynamic SCC Gaussians, then a subset of TPNs and TNNs are chosen to respond in the same way as in the Spectrally and Statistically Nonstationary regime.

#### SCC Gaussians: statistically stationarity without constraint on SDTEs

Following^9^, each simulated subject in the SCC Gaussian regime is a multivariate timeseries resulting from the projection of a 47 x 158 matrix of low-pass filtered white noise spectrally matched to 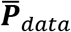 onto the eigenspace of 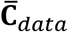 (see **Figure 7**; top middle).

#### Covariance-Dynamic SCC Gaussians: piecewise stationary, two distinct covariance regimes, no constraint on SDTEs (CD-SCC Gaussian)

This regime introduces explicit covariance nonstationarity. Each CD-SCC Gaussian subject starts as a 47 x 158 matrix consisting of low-pass filtered white noise spectrally matched to 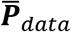, which is then divided into three windows, determined by a middle window of randomly chosen length between 40 and 60 TRs. The middle window is projected onto the eigenspace of 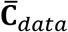, which has structure and is strongly connected, while the first and final windows are projected onto the eigenspace of ***C***_*weak*_, a covariance matrix reflecting very weak unstructured network connectivity (see **Figure 7**; bottom left).

#### SCC Gaussians with Noise: statistically stationary with a single spike randomly inserted into a small proportion of network timeseries (“Noisy SCC Gaussian”)

This regime introduces extremely sparse, high-amplitude noise to the SCC Gaussian setting. Each Noisy SCC Gaussian subject starts as an SCC Gaussian subject, i.e., as a 47 × 158 matrix consisting of low-pass filtered white noise spectrally matched to 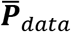 then covariance matched to 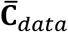. Of the 47 timeseries in this matrix, between *3* and 15 are selected at random to carry a single high frequency spike centered at some randomly selected timepoint. The entire multivariate timeseries contains between 3 and 15 of these noise artifacts, with at most one in any given univariate timeseries (see **Figure 7**; top right).

#### Spectrally and Statistically Nonstationary: explicitly nonstationary both statistically and epochally (“SS Nonstationary”)

This regime introduces simulated task-responsiveness to the SCC Gaussian setting. Each simulated SS Nonstationary subject starts as an SCC Gaussian subject (a 47 × 158 multivariate timeseries produced by subjecting white noise to spectral and covariance constraints exhibited by the real data). The 47 networks for each subject are divided into at most 17 task-positive networks (TPNs), at most 11 task-negative networks (TNNs), with the remaining 19 – 29 networks designated as nonresponders (NRs). Following the same procedure employed for the CD-SCC Gaussian regime, multivariate timeseries in the SS Nonstationary regime are divided into three windows determined by a middle window of randomly-chosen length between 40 and 60 TRs. The hypothetical task takes place during the middle window, in which (relative to the first and final window) the selected TPNs exhibit faster, higher amplitude behavior, the selected TNNs exhibit slower, lower amplitude behavior and the NRs exhibit no change in behavior (see **Figure 7**; bottom middle).

#### Covariance-Dynamic Spectrally and Statistically Nonstationary: explicitly nonstationary both statistically and epochally with two distinct covariance regimes (“CD-SS Nonstationary”)

This regime introduces explicit covariance nonstationarity to the SS Gaussian setting. Each simulated CD-SS Nonstationary subject starts as an SS Nonstationary subject (see immediately above). In this regime, however, the temporally task-responsive middle window is additionally subjected to explicitly different covariance constraints than the task-free first and final windows. Following the procedure from the CD-SCC Gaussian, the middle window of CD-SS Nonstationary subjects is projected onto the eigenspace of 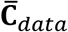, while the first and final windows are projected onto the eigenspace of ***C***_*weak*_ (see **Figure 7**; bottom right).

### 3.5. Kurtosis is Overly Sensitive to Noise Artifacts and Very Rarely Identifies Explicit Spectral, Statistical and Covariance Nonstationarities

As mentioned earlier, the utility of kurtosis as a measure of functionally relevant brain dynamics is mitigated by its highly-tuned sensitivity to spikes and outliers in the data. We saw this earlier in stylized univariate timeseries (see **Figure 5**). The issue is equally evident (see **Figure 8**) in more complex multivariate simulations involving different degrees of nonstationarity, exhibited in different ways. Employing m.v. kurtosis as a measure of FRBD implies that timeseries on which it assumes values large enough to provide significant evidence (*p* < 0.05) against the null hypothesis of stationary multivariate Gaussianity are those in which the features associated with FRBD are most markedly present. More specifically, these features increase monotonically with m.v. kurtosis. In **Figure 8** however, we see that that the only the simulation regime in which a non-negligible percentage (93%) of realizations is identified as exhibiting FRBD using m.v. kurtosis is the regime featuring a handful of spikes in an otherwise stationary 47 × 158 multivariate Gaussian. Only 5.3% of realizations from the explicitly covariance-dynamic regime are identified using the m.v. kurtosis metric as exhibiting FRBD, and for the other nonstationary regimes the percentage of realizations identified as exhibiting FRBD is less than a tenth of a percent. Similarly, the actual data recorded from subjects undergoing continual CASE-shifts exhibits no evidence by the m.v. kurtosis criterion of having arisen from a source in which functionally relevant brain dynamics are present. This clearly highlights the limitations of kurtosis as an indicator of FRBD. Elevated kurtosis indicates the sample contains too many points that are too extreme in magnitude under an assumption of stationary Gaussianity; it is less effective at capturing the dynamic changes in temporal behavior (including covariance) that are more plausible markers of FRBD and are richly present in most realizations of even stationary multivariate Gaussian processes.

**Figure 8.**
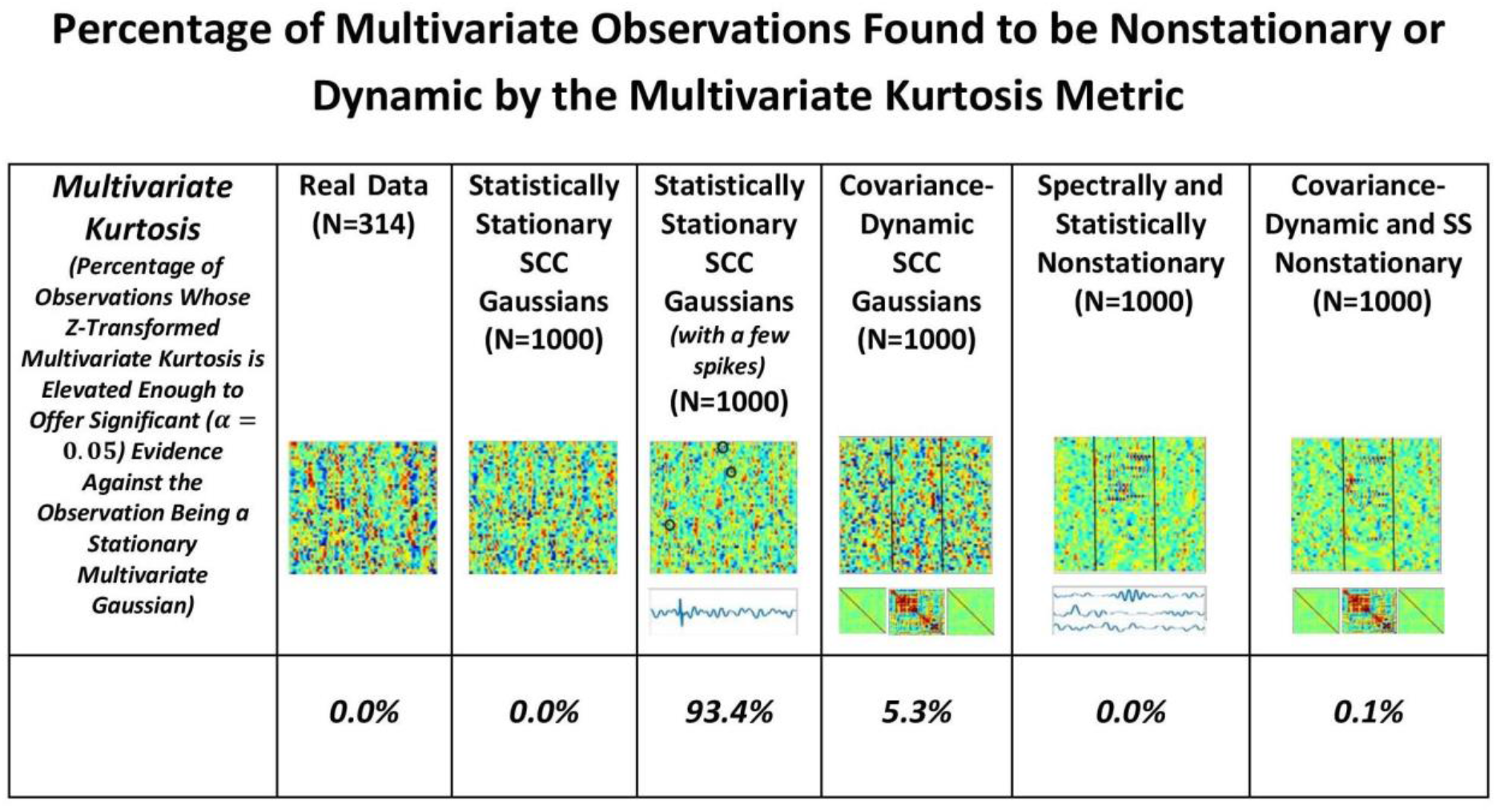
The percentage of multivariate timeseries from each indicated simulation regime (and the real rs-fMRI study on which the simulated data was modeled) that present significant evidence (*p* < 0.05) of having been generated by some process that is not a stationary multivariate Gaussian. The explicitly implemented spectral and statistical nonstationarities of the two SS Nonstationary regimes (columns 5 and 6) are effectively never found to exhibit significant evidence against being generated by stationary multivariate Gaussians. In the two explicitly covariance-nonstationary regimes (columns 4 and 6), one of which is also spectrally and statistically nonstationary variations (column 6), at most 5% of the 1000 simulated subjects – each of which exhibits the explicit nonstationarity – are identified as unlikely to have arisen from a stationary multivariate Gaussian process. Like the SCC Gaussian simulations (column 2), the SS Nonstationary simulations (columns 5 and 6) and the covariance-nonstationary simulations (columns 4 and 6), emprical observations from real subjects (column 1) in whom the phenomenon of interest (FRBD) is ubiquitous are not statistically distinguishable from realizations of a stationary multivariate Gaussian process. The only regime that multivariate kurtosis reliably distinguishes from realizations of a stationary multivariate Gaussian process is the case of SCC Gaussians in which a single high-amplitude, high-frequency spike is inserted into between 3 and 15 of the 47 univariate timeseries from the multivariate observation (column 3). This regime is basically just a lightly contaminated version of the SCC Gaussian regime (column 2) and of all of the simulation regimes exhibits the least evidence of functionally relevant brain dynamics. The behavior underlying upper-tail observations of multivariate kurtosis looks more like scan contamination than anything task-paradigm fMRI studies suggest would be strongly associated with FRBD.

Conversely, as shown in **Figure 9**, the proposed metric (defined in **Figure 4** and the Technical Supplement) responds in a more reasonable way to those features of real and simulated multivariate timeseries that have strong likelihood of reflecting FRBD vs. those that are simply aberrant in some other way. Results show that, in contrast to multivariate kurtosis, the lightly contaminated SCC Gaussians in terms of 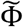 exhibit significantly less evidence of reflecting FRBD than either of the SS Nonstationary regimes (**Figure 9**, row 2, columns 5 and 6) and real data is statistically indistinguishable from the covariance-dynamic SCC Gaussian regime (**Figure 9**, row 1, column 4). So 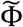 is putting regimes exhibiting different kinds of temporally epochal behavior—including the epochal behavior seen in statistically stationary Gaussian processes—in what seems a plausible ordering with respect to dynamism: SCC Gaussians ≼ Real Data ≈ Covariance-Dynamic SCC Gaussians ≼ Noise-Contaminated SCC Gaussians ≼ SS Nonstationary ≼ Covariance-Dynamic SS Nonstationary (where curly binary relations indicate ordinal evidence of potentially relevant multivariate epochal behavior as measured by 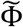 (see legend for more details).

**Figure 9.**
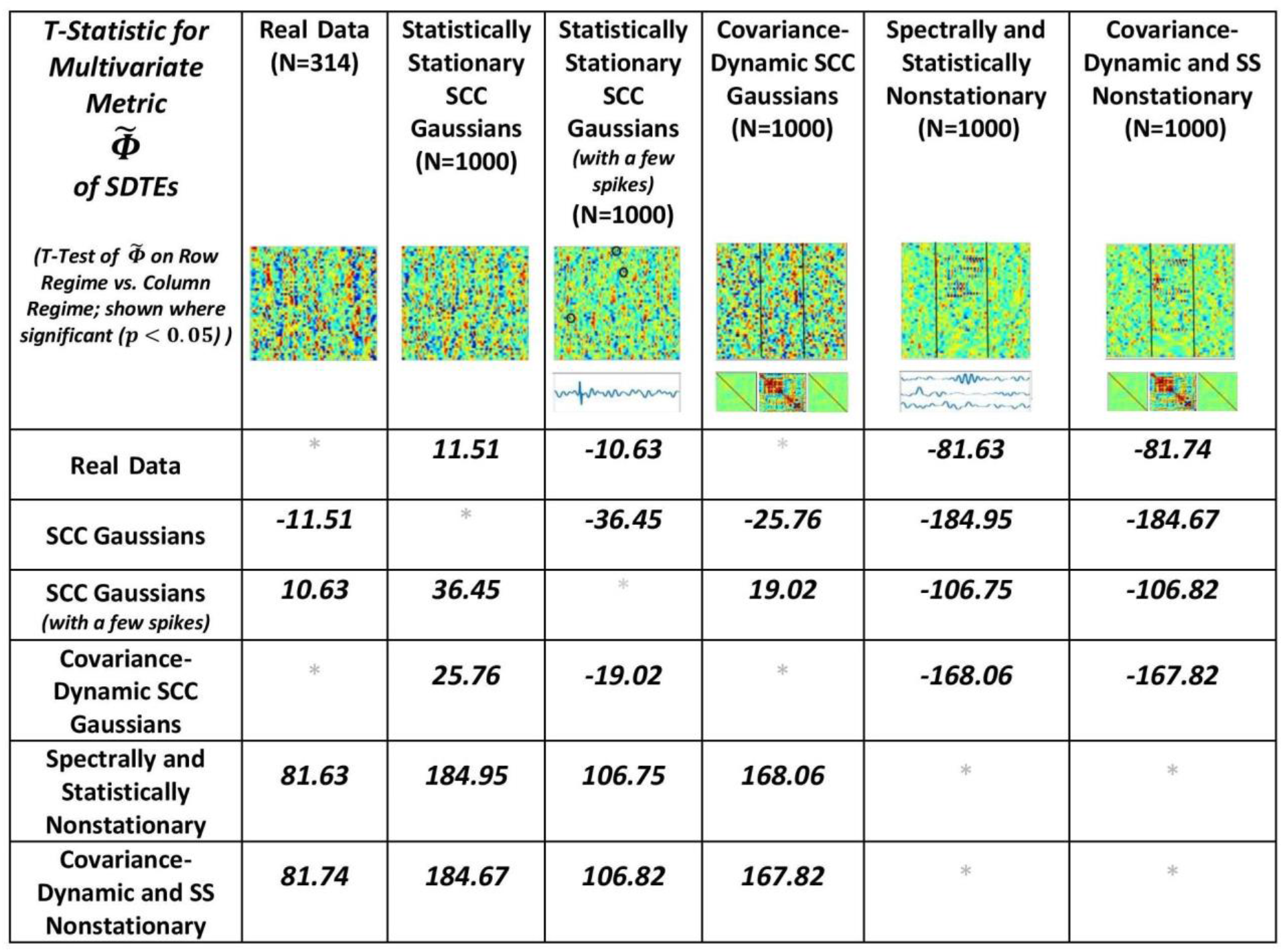
This table presents the t-statistic (where significant at the p<0.05 level) for pairwise T-tests of 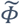 on the row regime vs. the column regime. From this standpoint, we see that the real network timecourses (row 1) exhibit significantly greater presence of SDTEs than the SCC Gaussians (column 2) modeled on them, significantly less evidence of SDTEs than the lightly contaminated “noisy” SCC Gaussians (column 3) and the explicitly SS Nonstationary Regimes (columns 5 and 6) and are statistically indistinguishable from the explicitly covariance-dynamic SCC Gaussians (column 4). Unlike what was found using multivariate kurtosis, the covariance-static SCC Gaussians (row 2) are in terms of ≼ significantly less dynamic than the covariance-dynamic SCC Gaussians (column 4) and both SS Nonstationary regimes (columns 5 and 6). Again, in contrast to multivariate kurtosis, the lightly contaminated SCC Gaussians (row 3) in terms of 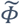 are significantly less dynamic than both SS Nonstationary regimes (columns 5 and 6). 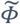 is putting regimes exhibiting different kinds of temporally epochal behavior – including the epochal behavior seen in statistically stationary Gaussian processes - in what seems a plausible ordering with respect to dynamism: SCC Gaussians ≼ Real Data ≈ Covariance-Dynamic SCC Gaussians ≼ Noise-Contaminated SCC Gaussians ≼ SS Nonstationary ≼ Covariance-Dynamic SS Nonstationary (where curly binary relations indicate ordinal evidence of potentially relevant multivariate epochal behavior as measured by 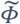).

This plausible ordering with respect to the t-statistic values from the analysis is much more meaningful for the φ metric than multivariate kurtosis. The spectra for the SCC Gaussians are more uniformly spread than the real data (please note that it is only the “average spectral power” that is matched to the real data) which is potentially one reason why the SCC Gaussians exhibit less evidence of dynamics than the real data. On the other hand, the multiple high frequency spikes add to temporal variation in the spectral content of the SCC Gaussians to which they are introduced, therefore resulting in false, elevated values of the multivariate metric in the modified (noisy) SCC Gaussian regime. Finally, both of the non-stationary classes are, by construction, explicitly nonstationary both statistically and epochally; hence, they are expected to exhibit greater evidence of dynamics. However less evidence of dynamics in real data than these two particular classes suggests that real data is not as non-stationary in nature as these explicitly nonstationary classes. In any case, preliminary advantages of using a raw measure based on these lines are clearly evident from the meaningful patterns as conveyed from these formal pairwise t-tests on metric values from the different regimes (**Figure 8**). Still, a lot more developmental and validation work is required to identify an exhaustive statistical measure most appropriate for the entire possible range/type of spectral and amplitudinal variations (or controlling ability for different classes of noise) in the resting state time-courses, and that validates as being statistically significant.

## 4. Choosing a Test Metric that Yields Different Distributional Tails for Null and Alternative Hypothesis

One key challenge in null model testing is choosing an appropriate combination of null model and test metric to allow for meaningful inferences. For a given null distribution to be truly null (for testing a given measure i.e. phenomenon of interest), the behavior of interest being evaluated must not be present generically (beyond a low probability level) within the null itself. Hence, the used test statistic must be sufficiently sensitive to the phenomenon of interest being tested for (as also discussed in^10^). The field of dynamic connectivity in particular has struggled with this as there are multiple threads of research proposing use of different models of dynamic behavior including covariance dynamics^1,18^ and oscillatory dynamics^19^. In this context, it is very important to keep any proposed null model within the narrow context within which it is able to reject a particular hypothesis about brain dynamics. For example, as seen earlier and also shown in^14^, a null model of oscillatory dynamics may well contain within it dynamically changing covariances, and, as such, it is not a particularly good null model for that particular scenario^10^. In addition to that, even models of covariance dynamics have been shown to be limited to a rather particular set of parameters, which we do not yet fully understand how to set for the human brain^8^. There is also evidence that the brain functions as a nonlinear dynamical system^20^. The result is that many of the null models that have been proposed are making strong assumptions about brain dynamics which, while having some justification, are not able to rule out dynamics of a different sort. We provide some additional discussion on this important point below.

### 4.1. Whole-Brain Windowed Connectivity States and Occupancies from Statistically Stationary and Explicitly Nonstationary Null Models Strongly Resemble Each Other

As we have discussed previously, creating a valid null model for rs-FRBD is difficult without explicitly understanding the properties that multivariate network timeseries might exhibit in response to complex CASE variations. One approach, suggested recently by^9^, employs a space of low-pass filtered multivariate white noise, spectrally matched to the average spectrum of empirical timecourse data and then projected onto the eigenspace of empirically observed mean network covariance. As discussed in Section 0, this is a space of timeseries explicitly modeled on real data recorded from a material in which the phenomenon of interest (i.e., functionally relevant brain dynamics) is continually present. It is therefore not a space in which univariate or multivariate temporal behavior plausibly sourced in CASE variations is vanishingly rare. Which is to say, it is not a useful null space for the identification of rs-FRBD. Moreover, it is a space of signals whose time-varying behavior is jointly determined by all simulation parameters and assumptions: the auxiliary spectral and covariance constraints as well as the Gaussianity assumption and the primary assumption of statistical stationarity. Above and beyond the problematic success of this simulation model in replicating empirical data recorded under circumstances in which the phenomenon of interest is uninterrupted and continual, this simulation regime’s value as a null model is further undermined by the unexamined role of auxiliary parameters in shaping key measures and distributions. This can be seen in **Figure 10**, rows 2 and 3, where we break the model’s core assumption of statistical stationarity, a property, arguably incorrectly, associated by the authors with an absence of FRBD, without discernibly disrupting either the clusters formed by short-timescale FNCs or the average cluster occupancy rates (see the Technical Supplement for a brief background on this approach).

**Figure 10.**
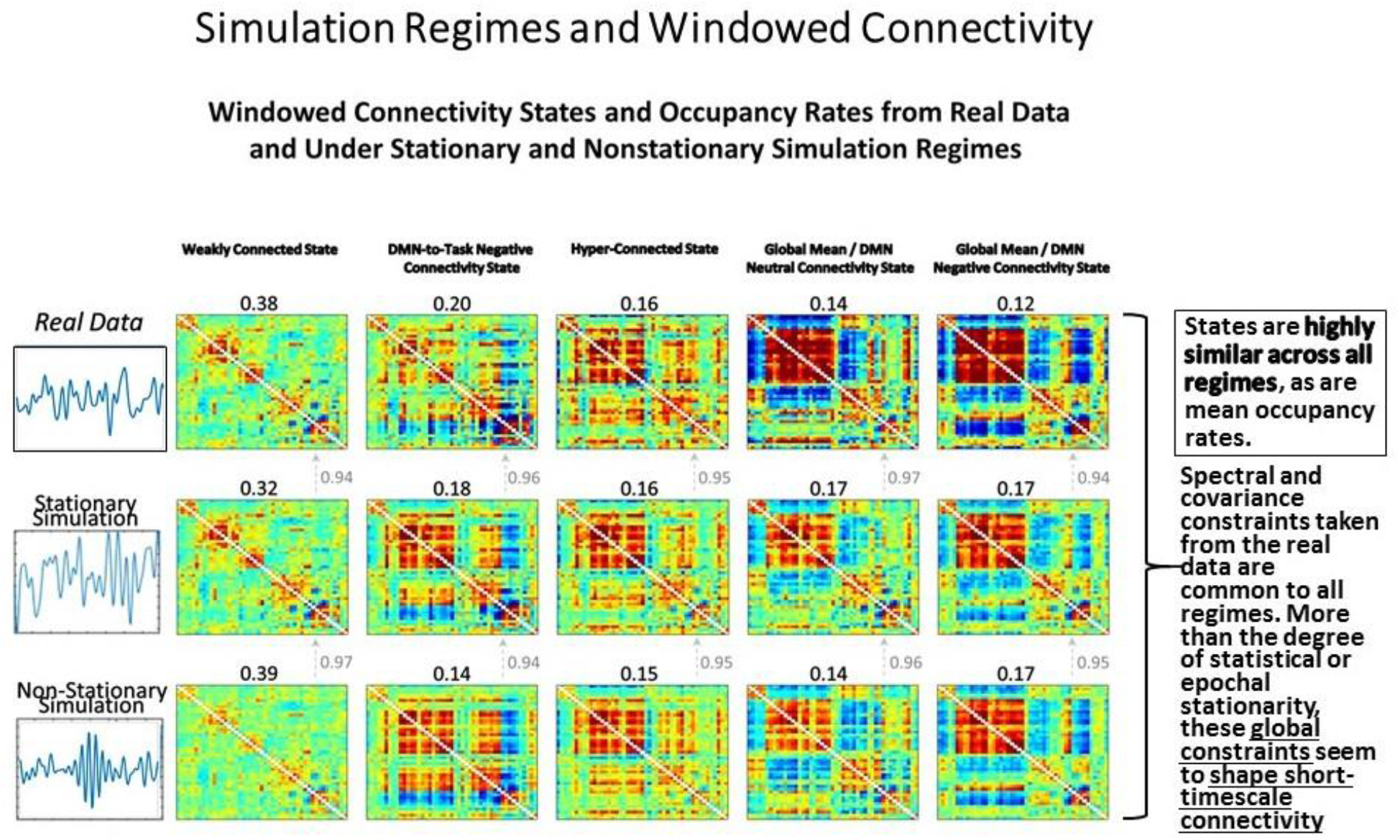
As we have pointed out previously, statistical stationarity, even multivariate stationary Gaussianity, can be richly inclusive of the types of time-varying multivariate behavior consistent with known brain responses to experimental tasks. Thus, the windowed connectivity states that a statistically stationary multivariate process moves through have every reason to resemble those of the real data upon which the process was tightly modeled (rows 1 and 2). However, it is also the case that short-timescale connectivity measurements from explicitly nonstationary processes (see “Statistically and Spectrally Nonstationary” regime in Section 3.4) subjected to spectral and global covariance constraints drawn from the real data cluster in the same way (row 3) as both the real data and the stationary Gaussian simulation modeled upon it. This suggests that the short-timescale connectivity states and occupancies are driven more by auxiliary constraints on mean spectrum and mean covariance than by whether the underlying process is statistically stationary. As we have shown, statistical stationarity does not preclude a multivariate signal from passing through connectivity states resembling those potentially arising from FRBD in real data. But more importantly it passes through the same connectivity states in the same way as explicitly nonstationary processes subjected to the same auxiliary constraints and as such is not particularly useful as a null condition for detection relevant brain dynamics.

The distributional tails of occupancy rates for each connectivity state in the SCC Gaussian and the SS Nonstationary regimes have significant overlap (see **Figure 11**) and as such this model can quite easily rule both for and against dynamic connectivity at the same time, an obvious flaw in the approach (i.e. evaluating statistical significance of measures of a phenomenon of interest generically present in a model chosen as the null as well as its alternative). This example illustrates the difficulties of building hypothesis-testing frameworks for phenomena whose distinguishing quantifiable characteristics are not well understood. If, in contrast to^9^, one realizes that the space within which one is working contains the very dynamics that one is trying to rule out (a point subsequently made by^14^) the conclusions that can be made are unconvincing and uninteresting.

**Figure 11.**
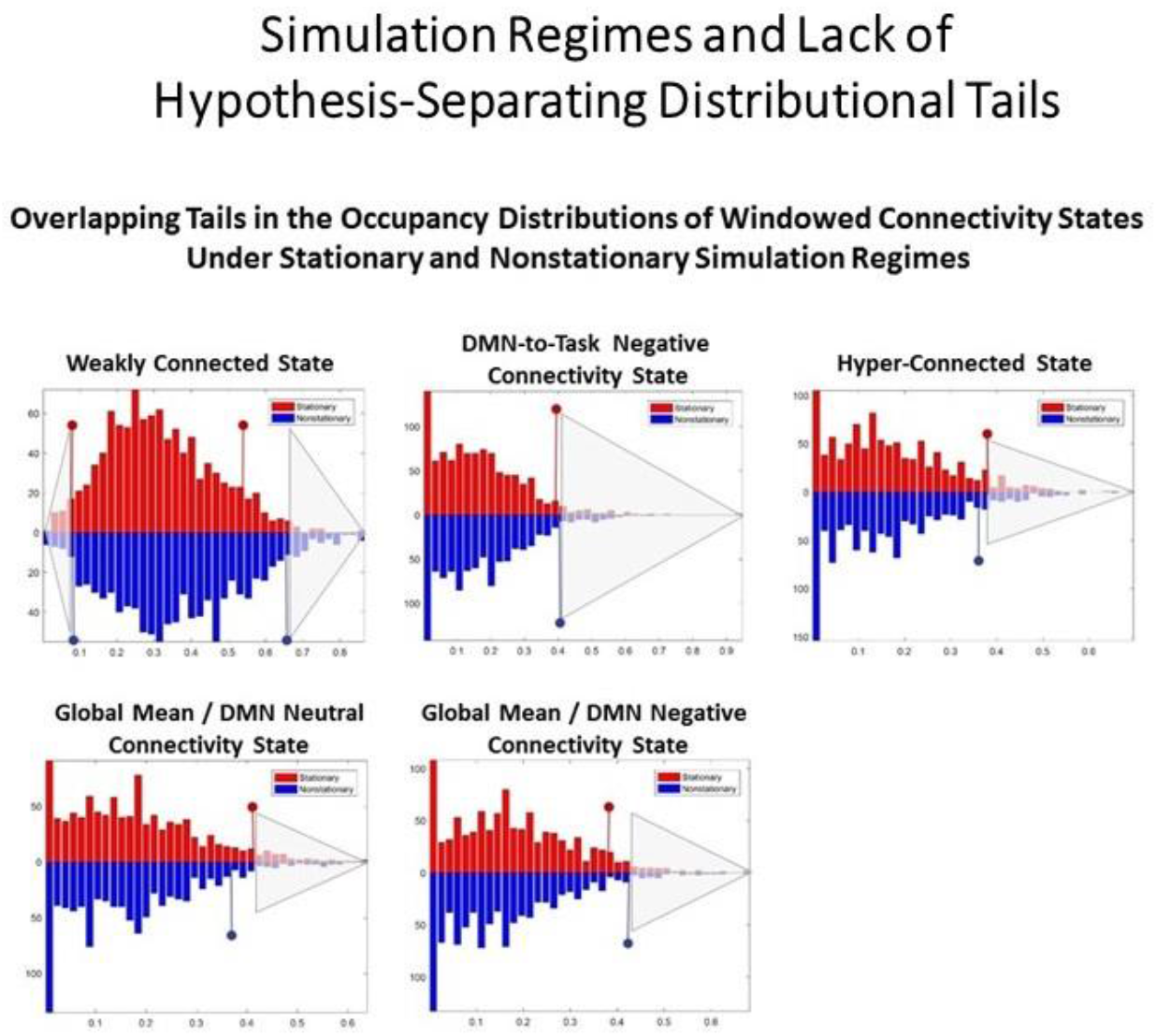
For a combination of null model and test metric to allow for meaningful inferences, the null model should exhibit evidence of the phenomenon being investigated (and believed to captured through the test metric) extremely rarely. SCC Gaussians arguably present evidence of FRBD as ubiquitously as the real fMRI data does. It is also not clear a priori that tail phenomena in SCC Gaussians should have an especially strong association to FRBD rather than, for example, to motion artifacts or measurement noise. Moreover, the measures evaluated on a null model are only helpful in identifying behavior of interest if the distributional tails of the measure do not overlap with those of the same measure evaluated on a model that violates assumptions of the original null hypothesis. The upper tails of the distributions of occupancy rates on the windowed connectivity states from SCC Gaussians (red) and the SS Nonstationary simulation regime (blue) have substantial overlap (semi-transparent grey triangles indicate the overlapping part of the tails for each state). Insufficient clarity on the unique, distinguishing features of the phenomenon being studied and of the models/measures being employed neutralizes the measure’s role in hypothesis testing as the same observation can then present significant evidence against multiple, mutually contradictory null hypotheses. In terms of occupancy rates of windowed connectivity states, we see here that a newly scanned subject who spends 65% of their time in state 2 exhibits significant evidence against one statistically stationary null hypothesis and also against an explicitly nonstationary null hypothesis.

## 5. Discussion

In this position paper, we have attempted to clarify and develop some of the important issues related to dynamic connectivity within the resting brain. Our focus has mostly centered around the use of metrics to detect possible dynamic behavior and also the creation of appropriate null models of functionally relevant brain dynamics, including but not limited to dynamic connectivity. We have discussed crucial limitations hindering some existing proposed models, demonstrated how a metric such as the one used in this paper could effectively detect potential properties of functionally relevant brain dynamics, and have, we hope, provided the context for a rich ongoing discussion of where the field should head from here. In the remainder of this paper, we highlight a few high-level questions that we hope make this point clear.

### 5.1. Are Functional Brain Dynamics Rare?

One of the key points we emphasized relates to the creation of a null simulation model. A useful null simulation model should be built on assumptions complementary to those that would apply if the phenomenon of interest were present, and then combined with a test statistic that is sufficiently sensitive to the null assumptions that it very rarely achieves extreme values when they are absent. Due to these concerns (as illustrated in the paper earlier), the null model in^9^ does not seem to be the right null model to evaluate functionally relevant brain dynamics with time-varying connectivity methods. A particular and carefully constructed null model was proposed in^10^, a study that evaluated statistical significance of variance as a test statistic to assess time-varying connectivity and emphasized importance of appropriate surrogate data testing in time-varying studies. The distributional tails of this test statistic on their static null simulation model matched the tails of variance computed on a wide range of explicitly dynamic variants of the model in which parameters were meaningfully changing with time. As also clearly pointed out by authors in this study^10^, such observations would simply suggest lack of sensitivity of this metric to separate out real fMRI data from stationary null model directly and would not suggest absence of brain dynamics in the fMRI data. Additionally, partially contrasting results have indeed been reported using the same null model (or null model retaining similar properties of the data) on the same metric^12^ or other time-varying measures factoring in this metric^13^. These studies have provided evidence that the stationary, linear and Gaussian model does not capture all the properties of the fMRI data; hence, the study of functional brain dynamics presents challenges to the utility of this entire type of modeling (to fit or study fMRI data). Moreover, we know that functional brain dynamics are constantly present in conscious human beings, so the phenomenon is anything but rare empirically.

### 5.2. Are Functional Brain Dynamics Separable from Nuisance Factors and Background Brain Rhythms?

The null model for a phenomenon necessarily produces, with very low probability, values of some test statistic that are consistent with the phenomenon which is being tested. This kind of model makes sense primarily when there is a very clear understanding of the range of values a particular test statistic will assume in the presence of the hypothesized phenomenon. So although in this short paper we are focusing primarily on the insufficiency of null models built on many realizations of some statistically stationary multivariate process modeled on empirical timecourse spectra and covariance, the larger problem is really that our present understanding cannot rule out the possibility of nontrivial intersection between signal features foreseeably connected with brain dynamics and those arising from nuisance factors and background brain rhythms. Even a spectrally pure signal, e.g., a single-frequency sinusoid, features amplitude changes that could be consistent with the ebb and flow of a network’s contributions to temporally varying CASE demands. Certainly pure sinusoids and other epochally stationary signals (whose phase-randomized stochastic analogues are statistically stationary) could provide evidence of network responsiveness in certain experimental task paradigms, e.g., those involving repetitive motor or sensory tasks. Since most temporal changes in a signal, including raw amplitude changes, could plausibly be correlated with some complex sequence of CASE conditions, the present state of knowledge makes it difficult to construct null models that can claim to yield, almost-exclusively, timeseries (multivariate or univariate) lacking features prospectively associated with brain dynamics.

This is true even when the null model is narrow. Indeed, it can be difficult to ensure that the test statistic being assessed does not have distributional tails roughly matched to the distributional tails of that statistic on a similarly narrow but explicitly “dynamic” model. In that case, we can easily conceive of examples where we are in the position of having an empirical measurement of the test statistic that simultaneously leads to rejection of one static null hypothesis and various related dynamic null hypotheses. In such a case, the desired test has been rendered essentially useless. Due to reasons seen earlier, the stationary null model is not the right null model to evaluate functionally relevant brain dynamics in future fMRI research. We expect functionally relevant temporal variations in brain activation to be constantly occurring throughout the experiment, and as the phenomenon of FRBD is better understood, improved null models will naturally emerge. We hope that the discussions herein are useful and promote thoughtful consideration of these important issues.

## 6. Conclusions

To summarize, serious and continuing investigation of dynamic multivariate brain activation patterns (including dynamic connectivity) is scientifically important and central to many core open questions in brain science. The time-varying measurements provided by BOLD fMRI currently play a vital supporting role in this overall project. We have discussed some of the limitations of existing null models and metrics for capturing dynamics, and provide initial evaluations of a new wavelet-based metric to demonstrate advantages of exploring more targeted measurements of time-series characteristics that form FRBD. While the evaluated metric appears to provide sensible results in a number of simulated scenarios, it still needs to be tuned and validated for an exhaustive range of spectral/amplitudinal variations and different noise classes. Moreover, a broader framework will ultimately be necessary to not only locate evidence of FRBD *per se* in univariate and multivariate brain data, but to also identify specific timepoints at which signals, signal-pairs and arbitrary signal *n*-tuples yield evidence that an underlying functional shift was underway. Finally, we urge caution in the development of null models in the context of dynamic connectivity. Especially for studies in which subjects are not engaging in a common, narrow experimental task, the relevant features, temporal and spatial/functional scales are not yet well understood. Specific well-defined questions about how particular signal features evolve on a range of spatial and temporal scales could produce more useful and testable hypotheses about how the brain signals we measure relate to high-level processes by which the brain organizes, directs and rotates through some of its central tasks: e.g., cognition, sense-making, generative thinking, memory-formation, memory-retrieval and emotion regulation, among others.

Finally, we would like to strongly emphasize that this paper is not an argument against the use of formal hypothesis testing in investigations of resting state dynamics. Nor is it an argument in favor of null spaces whose constituents have no recognizable relationship to actual brain data. In addition to tests for group differences or regression analyses (two cases in which there is always a well-defined null hypothesis), we are arguing in favor of narrower hypotheses that attach narrowly defined, non-ubiquitous neural/mental phenomena to a very narrow domain of signal characteristics. This will allow for null models that are, up to a limited set of signal properties, modeled on real fMRI data.

Following the choice of a narrow, non-ubiquitous target neural/mental phenomenon, we would advocate identifying some metric or test statistic *λ* on the data under which there is good reason to believe in the existence of an interval *I* (or region if *λ* is multidimensional) outside of which *λ* is both highly specific to and roughly monotonic in the presence (strength or abundance) of the target neural/mental phenomenon. This allows the construction of a useful null space built from simulated fMRI data that is as realistic as possible subject to the constraint that its elements induce values of *λ* outside of *I* at a rate consistent with the level of the target phenomenon an investigation is concerned with: very rarely if the target is present at levels of interest immediately beyond *I*, or less rarely if *λ* really is roughly monotonic and the target is only present at levels of interest as *λ* gets much further from *I*.

We would like to close by re-emphasizing our strong belief that hypothesis testing is crucial to advancing knowledge. The purpose of this paper is simply to highlight that in relatively young observational sciences the range of hypotheses that are rigorously testable at any given time might not extend to some questions of central importance to researchers. Most importantly, the set of rigorously testable hypotheses is constantly growing via ongoing vibrant interplay between more exploratory studies and studies that leverage the existing set of testable hypotheses for scientific gains.

## 7. Acknowledgement

This work was supported by National Institutes of Health (NIH) via a COBRE grant P20GM103472, R01 grants R01EB005846, 1R01EB006841, 1R01DA040487 and REB020407, and National Science Foundation (NSF) grant 1539067.

## Technical Supplement

### SCC Multivariate Gaussian Processes

Let be one 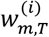 matrix of white Gaussian noise, 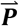 the average power spectrum for all subjects and all m network TCs in an actual resting state fMRI study, and ***C***_*m*_ the *m* × *m* population mean cross-network covariance. The set of timeseries one gets by inverse Fourier transforming a set of *T* normalized random complex coefficients 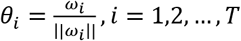 weighted according to a fixed template spectrum 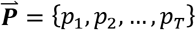, ie, the set:

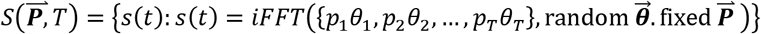

includes the discrete analogue (as we are working with computers on finite samples) of all length-*T* piecewise continuous functions with spectrum 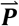. In particular, these timeseries exhaustively exhibit the full range of epochal behavior possible under the global spectral constraint 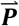. Since 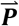 is the average spectrum of network TCs from a real rs-fMRI study, this ensures that the epochal signatures of CASE-driven rs-FRBD are replicated in 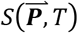. The set of *m*×*T* multivariate timeseries

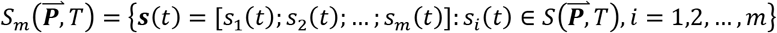

similarly exhaust the range of m-fold jointly realizable within-timeseries epochs subject to the duration *T* and spectral constraint 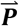. Finally, the set

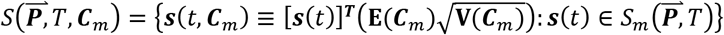

of projections

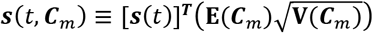

of 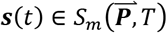 onto the eigenspace of ***C***_*n*_ is also a collection of Gaussian multivariate process with average spectrum 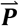 (where **E**(***C***_*m*_), **V**(***C***_*m*_) are respectively the eigenvectors and diagonal matrix of eigenvalues of ***C***_*m*_). The multivariate Gaussian timeseries in 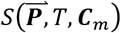 exhibit the joint-spectral and covariation epochs that can arise in *m*×*T* multivariate timeseries with average spectrum 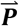 and whose full-duration covariance is brought into close approximation to ***C***_*m*_ by projection onto its eigenspace. No explicit constraints are imposed on shorter-timescale epochs of covariation within elements of 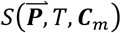; the set of possibilities is shaped primarily by ancillary constraints such as 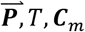 and the linear projection by which ***s***(*t*) is aligned toward ***C***_*m*_.

### Wavelet-Based Metric of Spectrally-Distinguishable Temporal Epochs

The metric 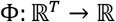 (still under active development) of within-timeseries spectrally distinguishable temporal epochs is computed on a univariate timeseries *S*(*t*), *t* ∈ {*t*_0_, *t*_1_, *t*_2_,…, *T*} as follows (see **Figure 5** in the main text):

1. Apply Matlab’s continuous wavelet transform (with the Morse wavelet) to *S*(*t*). This gives an *F*×*T* matrix *W*(*S*) of wavelet coefficient magnitudes. The rows of *W*(*S*) are timeseries of the power in each of the *F* frequencies. First rescale *W*(*S*) by the inverse of its mean, making its elements sum to *FT*.
2. The *T* × *T* symmetric matrix *D_Spec_*(*S*) of pairwise L1 distances between the columns of *W*(*S*) contains evidence of temporal variation in the core spectrum of *S. D_Spec_*(*S*) contains evidence of *epochal* spectral variation in *S*.
3. Finally, we set 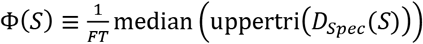 to be the median off-diagonal values of *D_Spec_*(*S*) rescaled by the average time-frequency power in *W*(*S*). Rescaling by the inverse summed power in *W*(*S*) keeps the value of Φ strictly bounded in [0,*T*].

The multivariate extension 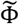 of Φ assumes that *W*(*S*) of size *F* × *T* has already been computed for some collection ***S*** of ***N*** length-*T* timeseries *S_i_, i* = 1,2,…,*N*. Let 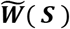 be the *NF*×*T* matrix of vertically concatenated *W*(*S_i_*)’s. Now 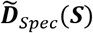 is the matrix of pairwise L1 distance between columns of 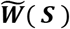 the concatenated spectra of the *S_i_*’s, and 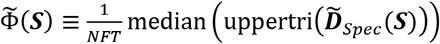 is the appropriately rescaled median off-diagonal value of L1 distances between the time-indexed concatenated spectra of the timeseries *S_i_, i* = 1,2,., *N*.

### Performance of the Wavelet-Based Metric for Small Changes in Spectral Content

The different univariate cases with small changes in amplitude and/or frequencies were evaluated and have been demonstrated in the figure below. Here are the details on the four toy time-series evaluated for kurtosis (*K*) and wavelet-based metric (Φ):

1. S_1_(t): Constant frequency and amplitude (used as reference);
2. S_2_(t): Constant frequency and slightly different amplitude;
3. S_3_(t): Slightly different frequency and constant amplitude; and
4. S_4_(t): Slightly different frequency and amplitude.

Higher values of the Φ metric were observed for the third and the fourth case, but not for the second case in comparison to the reference (first) case, which is understandable as the metric has been specifically designed to capture the temporal variations in the spectral content. Kurtosis did not show any significant change for any of these cases.

**Supplementary Figure 12:**
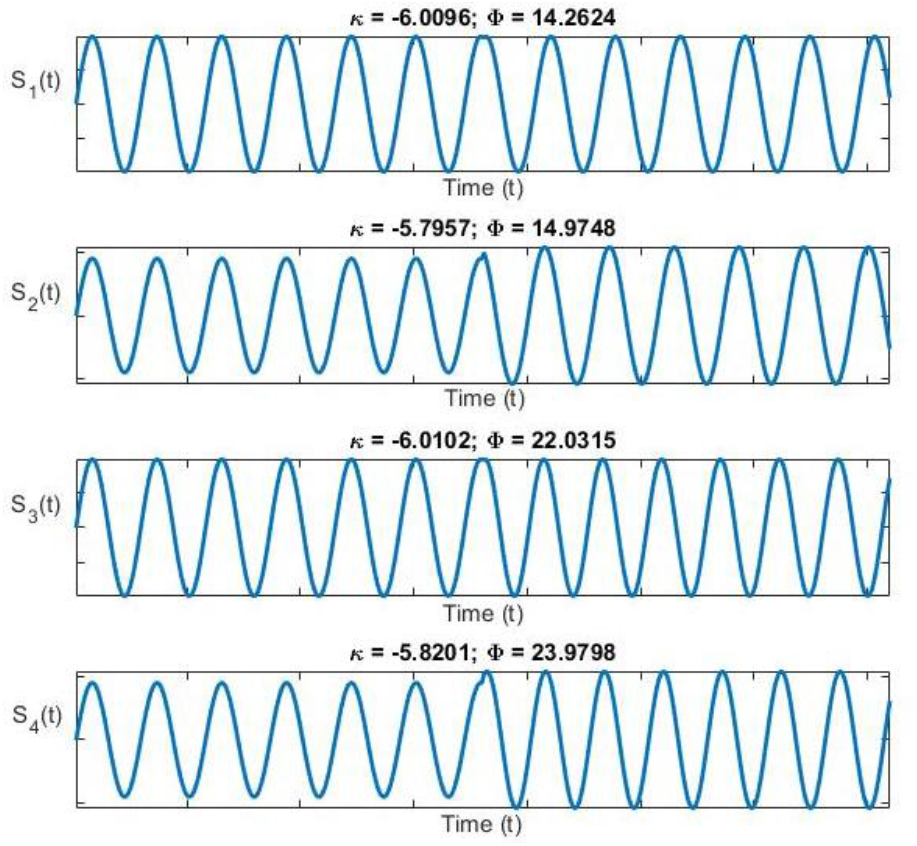
Elevated values of Φ (univariate wavelet-based metric) metric were observed for small temporal variations in spectral content. Kurtosis did not show significant change in any of these different scenarios.

The multivariate version of 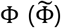 is computed from the median value of distances between concatenated time-indexed spectra; hence, temporal variation in spectral content of all constituent univariate time-series factors in the estimated value of this metric. Assuming epochs for all constituent univariate time-series to be varying slightly in frequency, the metric values could be expected to be lower than in cases where the temporal variation in spectral power is higher. We tested this case (slight changes in frequencies) and found that 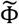 showed higher values in these cases as compared to the stationary reference case whereas multivariate kurtosis did not show any significant change as illustrated in the figure below. The toy example in Supplementary Figure 2 estimates both metrics in three different scenarios:

1. Each constituent time-series has a constant amplitude and frequency (used as reference for comparison);
2. Each constituent time-series has three equal intervals with constant amplitude but slightly different frequencies; and
3. Each constituent time-series has three equal intervals with slightly different amplitudes and different frequencies.

Scenarios 2 and 3 both show elevated values for 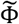 but similar values for multivariate kurtosis as compared to the reference scenario 1.

**Supplementary Figure 13:**
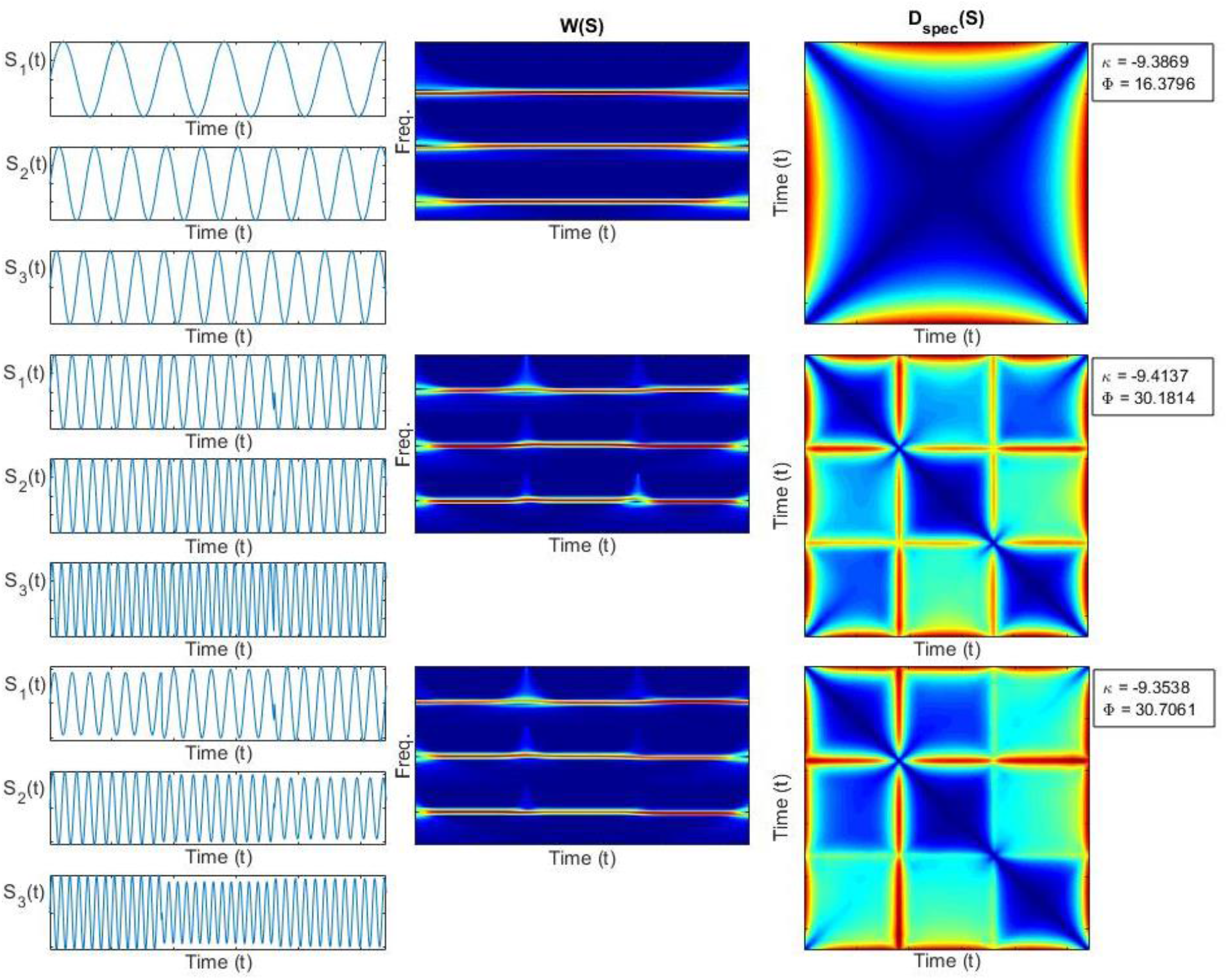
Elevated values of 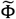 (multivariate wavelet-based metric) metric were observed for small temporal variations in spectral content. Similar values of multivariate kurtosis were reported in all scenarios.

### Empirical Data and Simulation Regimes

#### Real Data

We used previously published [1] network timecourse data from a large multisite clinical resting-state fMRI study. Preprocessing and network identification followed protocols detailed in [1] that we simply outline here. Resting state functional magnetic resonance imaging data (160 volumes of echo planar imaging BOLD fMRI, TR = 2 sec.) was collected from 163 healthy controls (117 males, 46 females; mean age 36.9) and 151 age and gender matched patients with schizophrenia (114 males, 37 females; mean age 37.8) during eyes closed condition at 7 different sites across United States. After standard preprocessing, the fMRI data from all subjects was decomposed using group ICA into 100 maximally spatially independent spatial maps (http://mialab.mrn.org/software) of which 47 were identified as functionally meaningful networks. The networks fell into seven broad categories: sub-cortical (SC), auditory (AUD), visual (VIS), sensorimotor (SM), cognitive control (CC), default mode network (DMN) and cerebellar (CB). Subject specific spatial maps and timecourses were obtained from the group level spatial maps via spatio-temporal regression. The timecourses were detrended, despiked and subjected to additional postprocessing steps detailed in [1].

These timecourses (314 subjects, 47 networks, 158 timepoints), further filtered for frequencies at most 0.08 Hz and then z-scored, are referred to below as “Real Data”. The average power at each frequency bin in [0.003,0.08] Hz for all network TCs for all subjects is denoted 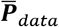. The average cross-network covariance matrix for all subjects is denoted 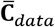.

Every simulation regime described below consists of 1000 subjects, each characterized by a set of 47, length-158 timeseries.

#### SCC Gaussians: statistically stationary without constraint on SDTEs

Following [2], each simulated subject in the SCC Gaussian regime is a multivariate timeseries resulting from the projection of a 47 × 158 matrix of low-pass filtered white noise spectrally matched to 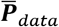 onto the eigenspace of 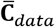 (**Figure 6** from the main text (top middle)).

#### Covariance-Dynamic SCC Gaussians: piecewise stationary, two distinct covariance regimes, no constraint on SDTEs (CD-SCC Gaussian)

Each simulated subject in the CD-SCC Gaussian regime starts as a set of 47 length-158 low-pass filtered white noise timeseries (organized in a 47 × 158 matrix) spectrally matched to 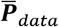. The multivariate timeseries is divided into three windows, determined by a middle window of randomly-chosen length between 40 and 60 TRs initially centered about the temporal midpoint *t_m_* = 79 then translated a random distance in [0,8] either forward or backward. This gives a middle window ([*t*_1_, *t*_2_],40 ≤ |*t*_2_ – *t*_1_ ≤ 60,41 ≤ *t*_1_ ≤ *t*_2_ ≤ 117) that is roughly central but differs in extent and in degree of centrality between subjects. This simulation regime employs two target covariance matrices: one, as usual, is the mean covariance 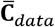 of the real data; the other 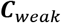 is modeled on 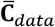 but with much weaker connectivity (except for preserved variances along the diagonal) and an additive layer of very low-magnitude Gaussian noise. For each subject, the random length middle window described above is projected onto the eigenspace of 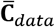, a modularly structured covariance matrix, while the first and final windows are each projected onto the eigenspace of ***C***_*weak*_. The resulting multivariate timeseries are explicitly undergoing a covariance shift as they enter and leave a 40-60TR window spanning the temporal midpoint of the scan (**Figure 6** from the main text (bottom left)).

#### SCC Gaussians with Noise: statistically stationary with a single spike randomly inserted into a small proportion of network timeseries (“Noisy SCC Gaussian”)

Each simulated subject in the Noisy SCC Gaussian regime starts as an SCC Gaussian subject (a 47 × 158 multivariate timeseries produced by subjecting white noise to spectral and covariance constraints exhibited by the real data). Of the 47 timeseries in this matrix, between 3 and 15 are selected at random to carry a single high frequency spike. This spike is a high-frequency, high-amplitude artifact centered at some randomly chosen timepoint in each of the selected networks The entire multivariate timeseries contains between 3 and 15 of these noise artifacts, with at most one appearing in any given univariate timeseries (**Figure 6** from the main text (top right)).

#### Spectrally and Statistically Nonstationary: explicitly nonstationary both statistically and epochally (“SS Nonstationary”)

Each simulated subject in the Nonstationary regime starts as a SCC Gaussian subject (a 47 × 158 multivariate timeseries produced by subjecting white noise to spectral and covariance constraints exhibited by the real data). We designate 22 of the 47 simulated networks, (with row-indices corresponding to those of auditory, visual, sensorimotor and select cognitive control networks in ***C***_*dat*_) as task-positive (TPNs). Another of 15 of the remaining 25 networks (with row-indices corresponding to default mode networks and select subcortical networks in ***C***_*dat*_) as task-negative (TNNs). For each subject, a randomly selected 50%−75% of the TPNs and 50–75% of the TNNs are selected to exhibit stylized responsiveness to a hypothetical stimulus, leading each simulated subject to have at least 19 and most 29 responders among their 47 networks. Following the same procedure employed for the CD-SCC Gaussian regime, multivariate timeseries in the Nonstationary regime are divided into three windows whose endpoints are determined by a middle window of randomly-chosen length between 40 and 60 TRs initially centered about the temporal midpoint *t_m_* = 79, then translated a random distance in [0,8] either forward or backward. As mentioned above, this gives a middle window ([*t*_1_,*t*_2_],40 ≤ |*t*_2_ – *t*_1_| ≤ 60,41 ≤ *t*_1_ ≤ *t*_2_ ≤ 117) that is roughly central but differs in extent and in degree of centrality between subjects. In those networks selected as responding TPNs, the middle window is filtered for frequency content in [0.06,0.08] Hz, then rescaled to have amplitude slightly higher than the first and last windows. In responding TNNs, the middle window is filtered for frequency content in [0.006,0.05] Hz and rescaled to have amplitude slightly lower than the first and last windows. So TPNs get faster and stronger in response to the hypothetical task or stimulus, while TNNs go into a slower shallower activation regime. The 19–29 nonresponding networks remain as they were following the initial spectral and covariance-matching steps. This yields a 47 × 158 matrix in which 11–17 rows contain TPN timeseries, each with a fast high-amplitude middle window, 7–11 rows contain TNN timeseries, each with slow shallow middle window and 19–29 low-pass filtered (non-windowed) SCC Gaussians unchanged after the initial spectral and covariance-matching step. This set of multivariate timeseries is explicitly spectrally and statistically nonstationary (**Figure 6** from the main text (bottom middle)).

#### Covariance-Dynamic Spectrally and Statistically Nonstationary: explicitly nonstationary both statistically and epochally with two distinct covariance regimes (“CD-SS Nonstationary”)

Each simulated subject in the CD-Nonstationary regime starts as a Nonstationary subject as defined immediately above. However, following the CD-SCC Gaussian regime, this simulation regime employs the two, distinct target covariance matrices detailed above in the description of the CD-SCC Gaussian regime. For each subject, the random length middle window in which a subset of networks is spectrally perturbed (as detailed in the section immediately preceding) is projected onto the eigenspace of the modularly structured covariance matrix 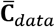 while the first and final windows are each projected onto the eigenspace of ***C***_*weak*_. The resulting multivariate timeseries are explicitly undergoing spectral, amplitude *and* covariance shifts as they enter and leave a 40-60TR window spanning the temporal midpoint of the scan (**Figure 6** from the main text (bottom right)).

#### Univariate and Multivariate Kurtosis

Univariate kurtosis, is the fourth statistical moment, 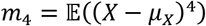 of a random variable *X*, rescaled by the variance 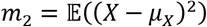 squared. For a normal random variable 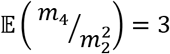, so *excess univariate kurtosis* indicative of super-Gaussianity (unusually heavy tails) is given by 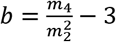. An unbiased estimator of excess univariate kurtosis [3] is:

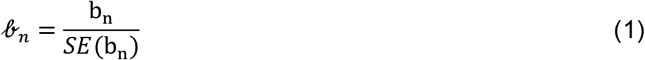

where

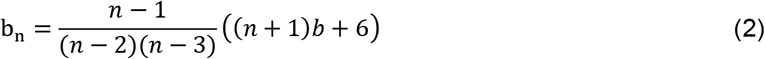

and

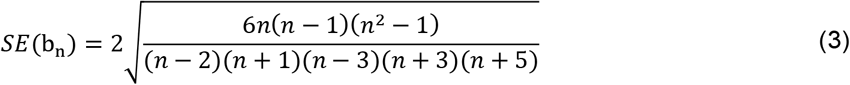

Multivariate kurtosis is a generalization of univariate kurtosis introduced by [4]. For a given length-*n* multivariate process consisting of *p* univariate timeseries, Maria’s multivariate kurtosis is defined as

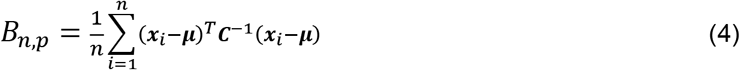

where

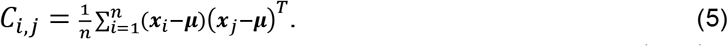

For a multivariate Gaussian process consisting of *p* univariate timeseries of length *n* → ∞, 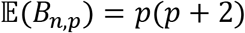. Excess multivariate kurtosis, *B_n,p_* – *p*(*p* + 2) is evidence of multivariate super-Gaussianity. An unbiased estimator [5] of excess *B_n,p_* is:

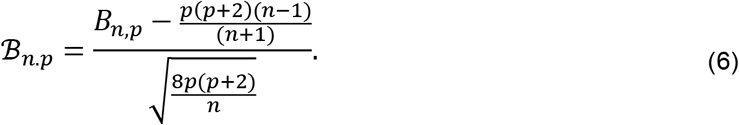

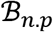 has a standard normal distribution 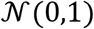 when the underlying process is actually multivariate Gaussian, so the upper tail represents strong evidence against the null hypothesis that all observations all arose from fixed Gaussian distributions in each dimension.

#### Sliding Window Dynamic Connectivity

Short-timescale network or region of interest (ROI) connectivity estimates evaluated on successive sliding windows through activation timecourses are a commonly employed [1, 6–9] vehicle through which to investigate so-called dynamic connectivity in resting-state fMRI. The general idea is straightforward: starting from a set of *R* length-*T* network or ROI timecourses that emerges from a standard pre-preprocessing pipeleine for rs-fMRI, slide a window of fixed length *L* vertically through each subject’s *T*×*R* multivariate timeseries, advancing stepwise by some increment *l* until the whole timeseries is exhausted. The window can be rectangular or have tapered edges; the pipeline that selects networks or ROIs and processes the resulting timeseries is generally unaltered relative to the non-windowed setting, and the metric of connectivity – often but not always correlation or covariance – is typically not unaltered from the non-windowed setting, in part to allow for comparisons between static (i.e., scan-length) findings and windowed (short time-scale) findings. There is considerable debate [10–13] regarding appropriate window-length subject for example, to the spectral content of the signal and other considerations. However our choice of 44s (similar to window duration as used in [1] on the same fBIRN phase 3 dataset) falls within previously recommended ranges. In background, [14] formally demonstrated use of the proposed lower limit of window length using the (inverse of minimum frequency) thumb rule as originally proposed in [12] to be overly conservative especially in moderate SNR conditions (i.e. relatively much shorter windows than as suggested by the thumb rule can be used to capture the fluctuations in time-varying connectivity). Furthermore, in their simulations, the authors in [10] indeed demonstrate that the maximum probability of detection peaks around 50 seconds. Similarly, [15] found peaks of significance of window lengths in the 40-60 seconds range. Moreover, there are several studies that corroborate that varying the window length parameter over a range beyond a certain safety limit did not change the overall observed dynamics [9, 16–19]. In summary, window size does have a substantial impact on the time-varying FC estimates as pointed out in recent papers [10–13], but recent work seems to be convergent around the 40 to 60 seconds range. Furthermore, due to the number of connectivity measurements this approach generates per subject, it is common to attempt to summarize the short-timescale connectivity patterns in the study by clustering, using fc-means, the entire set of windowed observations, leading to some collection of *k* summary connectivity states. These *k* connectivity states summarizing transient connectivity patterns in the entire population yield easily computable information (e.g., occupancy rates, mean dwell times and transition probabilities) about subject-level time-varying connectivity.

As indicated above, in the section about empirical data and simulation regimes, we used previously published [1] network timecourse data from a large multisite clinical resting-state fMRI study as the empirical basis for simulation models. The original study filtered timecourses for spectral content under [0.003,0.125] Hz, but in keeping with [2], in this paper we filtered for content [0.003,0.08] Hz. Otherwise our initial pipeline leading up to windowing was identical to that published in [1] (and outlined above in the section on empirical data and simulation regimes). Again following the published study [1], our windows had length 22 TR and were advanced by 1 TR at each step, leading to a total of 136 windows per subject. Although [1] used Gaussian tapering, here we employed rectangular windows. There was no discernible difference between the two approaches, so the simpler approach was utilized. Connectivity between networks on each window is measured as pairwise correlation between the windowed network timecourses. In the original study, the elbow criterion suggested *k* = 5 clusters for this data, a choice that we retain here. For consistency and comparability between empirical data and the various simulation regimes, all simulation regimes were windowed and clustered (using *k* = 5) with the same protocols and parameters as the empirical data.

## Footnotes

1 The term “epoch” has been used in this paper to refer to a duration, time period, and not necessarily of repetitive nature.

